# A likelihood-based deconvolution of bulk gene expression data using single-cell references

**DOI:** 10.1101/2020.10.01.322867

**Authors:** Dan D. Erdmann-Pham, Jonathan Fischer, Justin Hong, Yun S. Song

**Affiliations:** Department of Mathematics, University of California, Berkeley, California 94720; Department of Statistics, University of California, Berkeley, California 94720; Computer Science Division, University of California, Berkeley, California 94720; Department of Biostatistics, University of Florida, Gainesville, Florida 32611; Chan Zuckerberg Biohub, San Francisco, California 94158

## Abstract

Direct comparison of bulk gene expression profiles is complicated by distinct cell type mixtures in each sample which obscure whether observed differences are actually due to changes in expression levels themselves or simply due to differing cell type compositions. Single-cell technology has made it possible to measure gene expression in individual cells, achieving higher resolution at the expense of increased noise. If carefully incorporated, such single-cell data can be used to deconvolve bulk samples to yield accurate estimates of the true cell type proportions, thus enabling one to disentangle the effects of differential expression and cell type mixtures. Here, we propose a generative model and a likelihood-based inference method that uses asymptotic statistical theory and a novel optimization procedure to perform deconvolution of bulk RNA-seq data to produce accurate cell type proportion estimates. We demonstrate the effectiveness of our method, called RNA-Sieve, across a diverse array of scenarios involving real data and discuss extensions made uniquely possible by our probabilistic framework, including a demonstration of well-calibrated confidence intervals.

## Introduction

Bulk RNA sequencing (RNA-seq) has proven a useful tool to investigate transcriptomic variation across organs/tissues, individuals, and various other biological conditions (Melé et al., 2015; Sudmant et al., 2015). Despite many successes, this technology’s full potential is inherently limited because each experiment measures the average gene expression among a large group of cells, the composition of which is unknown. Thus, despite the reduction in technical and biological variability attained by averaging, bulk experiments are potentially confounded by cell type proportions when considering heterogeneous cell mixtures (Lowe and Rakyan, 2014; Shiwa et al., 2016). Such confounding impedes the direct comparison of samples, possibly leading to the spurious or missed inference of biologically relevant genes when attempting to identify clinically important differences. Moreover, cell type compositions are often independently informative of biological processes including organ function (Cabrera et al., 2006; Hagenauer et al., 2018; Kalisky et al., 2013; Yu and He, 2017) and development (Hagenauer et al., 2018; Hu et al., 2017). For example, cell type infiltration has been found to correlate with disease progression, disease status, and complex phenomena such as aging (Funada et al., 2003; Bense et al., 2017; Zhou et al., 2019; Bremnes et al., 2016; Stout et al., 2017). Unlike bulk experiments, single-cell technologies allow us to query the transcriptome at the resolution of individual cells. Resulting analyses often seek to characterize the heterogeneity within, or the differences between, specified cell types (Saliba et al., 2014). By isolating the expression patterns of each measured cell type, single-cell gene expression data can provide a reference to aid the inference of the cell type compositions of bulk samples; this process is known as deconvolution.

Computational rather than experimental estimation of cell type compositions is attractive for several reasons. Single-cell experiments are more expensive than their bulk counterparts and require heightened technical expertise to perform, often rendering the large-scale generation of singlecell gene expression data infeasible (Goldman et al., 2019). Furthermore, even when performed correctly, many protocols fail to capture cell types in an unbiased fashion, meaning empirical cell type proportions often are not reliable estimators of true organ/tissue compositions (Trapnell, 2015). Finally, deconvolution can be applied to the deep compendium of available bulk RNA-seq data to refine earlier analyses and probe previously unanswerable or heretofore unformulated questions. Consequently, the computational deconvolution problem has become a topic of intense methodological research (as detailed in Avila Cobos et al. (2018)). The problem may be represented mathematically as

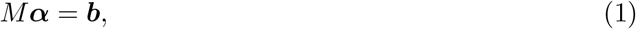

where *M* is a gene-by-cell type matrix of cell type-specific gene expression averages, ***α*** a vector of cell type mixing proportions, and ***b*** a vector of gene expression values in a bulk RNA-seq experiment. Depending on which of *M*, ***α***, and ***b*** are measured, different approaches are appropriate. We focus on the case in which both *M* and ***b*** have been observed, albeit noisily, and it remains to infer ***α***; this is known as supervised deconvolution. Early approaches frequently utilized pre-defined marker genes for well-studied cell types, restricting their applicability. More recent methods formulate the problem as a regression task to be solved by variants of non-negative least squares (e.g., MuSiC (Wang et al., 2019), DWLS (Tsoucas et al., 2019), SCDC (Dong et al., 2020), and Bisque (Jew et al., 2020)) or with more sophisticated machine learning techniques (such as CIBERSORTx (Newman et al., 2019) and Scaden (Menden et al., 2020)). Though each paradigm presents its own strengths, both fail to replicate the benefits of explicit generative modeling. The result is algorithms which may perform well but lack the flexibility to extend beyond the estimation of cell type proportions.

In this paper, we propose a new method, RNA-Sieve, which employs asymptotic theory and a novel optimization procedure to solve a probabalistic model of deconvolution via maximum likelihood estimation. We demonstrate its highly capable performance across a diverse array of scenarios, including different organs/tissues, cell types, and practical challenges. We then highlight newly opened avenues for continued development made feasible by our generative framework, including confidence regions and general hypothesis tests.

## Results

### Method Overview

Although a single run of bulk RNA-seq produces only a solitary gene expression vector, myriad cells contribute to this measurement. The obtained profile is hence a composite snapshot of the gene expression levels of numerous individual, putatively independent cells. When coupled with an assumption that cells of the same type behave similarly, this large number of cells permits the application of the central limit theorem (CLT) and the wealth of normal distribution approximations it implies. Conveniently, the marginal distribution for gene expression of an arbitrary cell in the bulk sample is a straightforward mixture distribution (see Equation (2) in Methods). The resulting CLT-derived likelihood consequently depends only on the means and variances of gene expression for each cell type and the respective cell type proportions in the bulk sample, the latter of which we make our goal to infer computationally. To estimate the requisite cell type-specific moments, RNA-Sieve uses gene expression measurements from scRNA-seq experiments. We further model the estimation error of the computed moments by once again invoking the CLT, and the combination of these two approximations yields a full, composite likelihood built using normal distributions. We subsequently infer cell type proportions via a custom-made maximum likelihood estimation procedure. Several features of our algorithm ensure accurate and robust results. Our alternating optimization scheme is split into two components to better avoid sub-optimal local minima, with a final projection step handling flat extrema to avoid slow convergence. We also incorporate a gene filtering procedure explicitly devised to improve cross-protocol stability, a crucial concern given that single-cell and bulk experiments will always be performed with different technological platforms. Our algorithm can also perform joint deconvolutions, leveraging multiple samples to produce more reliable estimates while parallelizing much of the optimization. In this setting, each included bulk sample improves the denoising of the single-cell reference regardless of its mixture proportions, leading to improved statistical performance. Finally, we wish to emphasize that our likelihood-based model allows us to pursue extensions which are infeasible using prior approaches. A notable example includes confidence regions for estimates (see Extension to Confidence Regions), among others (see Discussion). Full mathematical and computational details are presented in Methods, and a schematic is displayed in Figure 1.

**Figure 1:**
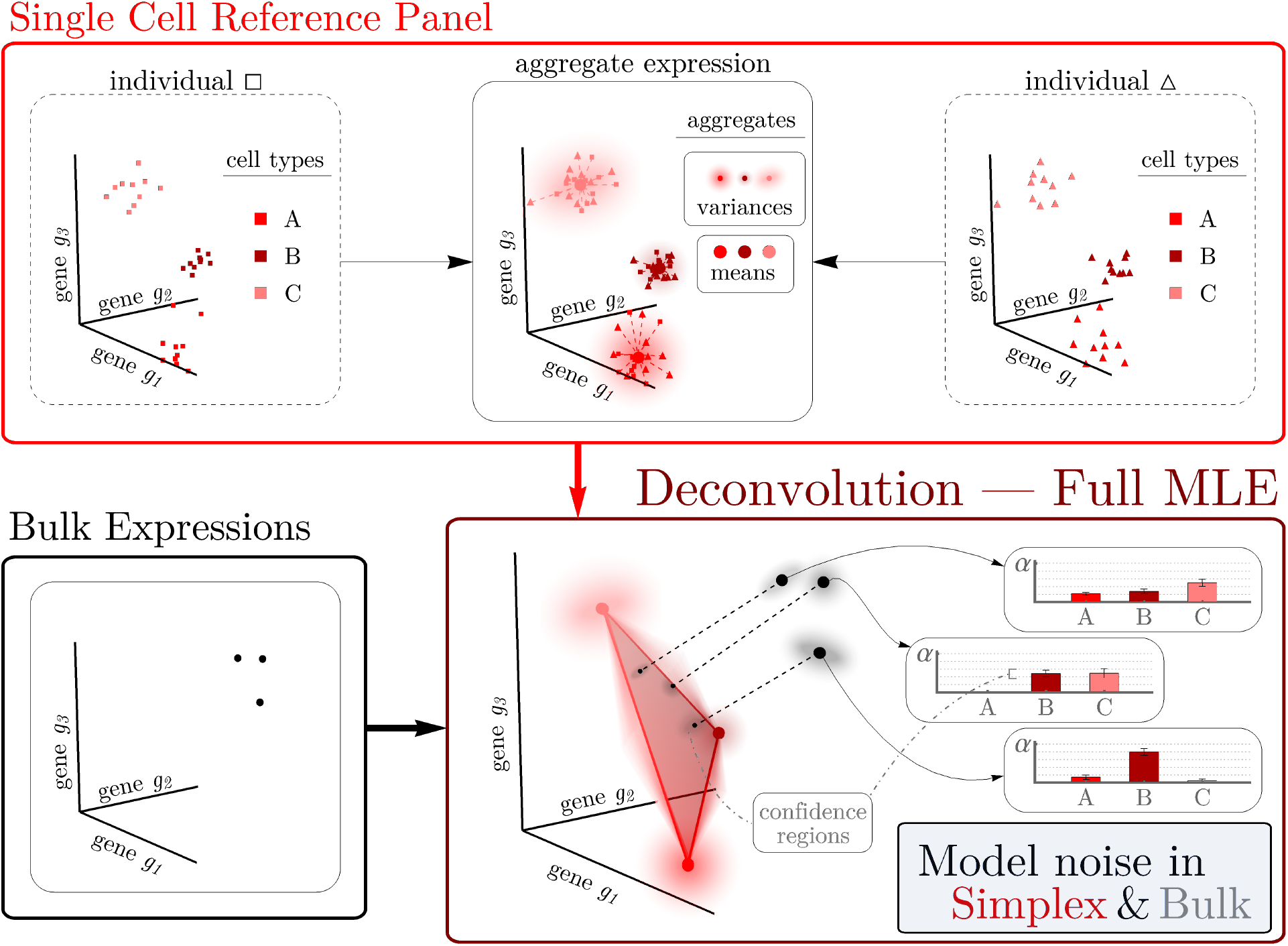
The RNA-Sieve pipeline. After applying a filtering procedure to scRNA-seq data, RNA-Sieve builds reference matrices for the mean and variance of expression for each gene across cell types. Using these estimates and bulk RNA-seq data, it performs joint deconvolution via maximum likelihood estimation by expressly modeling noise both in the reference and bulk data, yielding cell type proportion estimates and confidence regions for each sample.

### Performance in Pseudobulk Experiments

To establish RNA-Sieve’s effectiveness, we performed *in silico* experiments using scRNA-seq data from the *Tabula Muris Senis* Consortium (Tabula Muris Consortium et al., 2020). In these experiments we built “pseudobulks” by aggregating reads from labeled cells in known proportions to use for deconvolution. We considered thirteen organs with between two and eleven cell types per organ. Moreover, counts generated via both the Smart-Seq2 and 10x Chromium protocols are available for each organ, enabling convenient cross-protocol comparisons. These are particularly important given that bulk and single-cell RNA-seq samples are always processed using different techniques. To evaluate RNA-Sieve, we compared its performance to that of six recently published methods as well as non-negative least squares (NNLS). Performance was assessed for each organ by computing the *L*_1_ distance (absolute difference) between inferred and true proportions and dividing by the number of cell types present. Further details are provided in Benchmarking procedures. We found that RNA-Sieve produced the smallest mean error in both possible reference/bulk configurations (Figure 2 and Table 1; full results in Table S1). To better understand performance, we also visualized errors when aggregating by organ (that is, the column-wise distributions of the checkerboard plots in Figure 2, see Figure S1). This demonstrated that RNA-Sieve performed at least as well as all competitors, and often notably better, in nearly all organs. Our strong performance across organs regardless of the number of cell types or similarities among them suggests that RNA-Sieve is versatile over a range of scenarios. Finally, we directly compared each method’s errors to those of RNA-Sieve on the same deconvolution tasks (given by the row-wise distributions of the checkerboards in Figure 2, see Figure S2). In each case, RNA-Sieve produced smaller errors than the other methods a majority of the time.

**Table 1:** Summary of deconvolution errors for each considered method in pseduobulk experiments. Errors were computed as the *L*_1_ distance (in %) between the inferred and true proportions averaged over the number of present cell types per organ. Single-cell RNA-seq data for the references and pseudobulks were taken from the *Tabula Muris Senis* experiment. The mean, median, and interquartile range are displayed for the results in thirteen different organs; see Benchmarking procedures for additional details.

| (a) Smart-Seq2 reference and 10x Chromium pseudobulk. |  |  |  |  |  |  |  |  |
| --- | --- | --- | --- | --- | --- | --- | --- | --- |
|  | RNA-Sieve | CIBERSORTx | Scaden | SCDC | MuSiC | DWLS | Bisque | NNLS |
| Mean | **6.9** | 8.6 | 8.6 | 8.7 | 10.1 | 11.2 | 12.7 | 30.5 |
| Median | **6.1** | 7.1 | 7.2 | 8.3 | 10.6 | 7.2 | 10.1 | 31.0 |
| IQR | 7.7 | 3.4 | 4.1 | 7.9 | 6.0 | 6.9 | 5.2 | 15.3 |

| (b) 10x Chromium reference and Smart-Seq2 pseudobulk. |  |  |  |  |  |  |  |  |
| --- | --- | --- | --- | --- | --- | --- | --- | --- |
|  | RNA-Sieve | CIBERSORTx | DWLS | Scaden | Bisque | SCDC | MuSiC | NNLS |
| Mean | **6.7** | 7.4 | 7.6 | 10.5 | 12.5 | 18.1 | 18.7 | 26.4 |
| Median | 8.2 | 6.2 | **5.4** | 10.5 | 11.0 | 15.7 | 15.7 | 16.4 |
| IQR | 9.9 | 7.8 | 6.9 | 4.6 | 8.9 | 12.1 | 4.6 | 17.0 |

**Figure 2:**
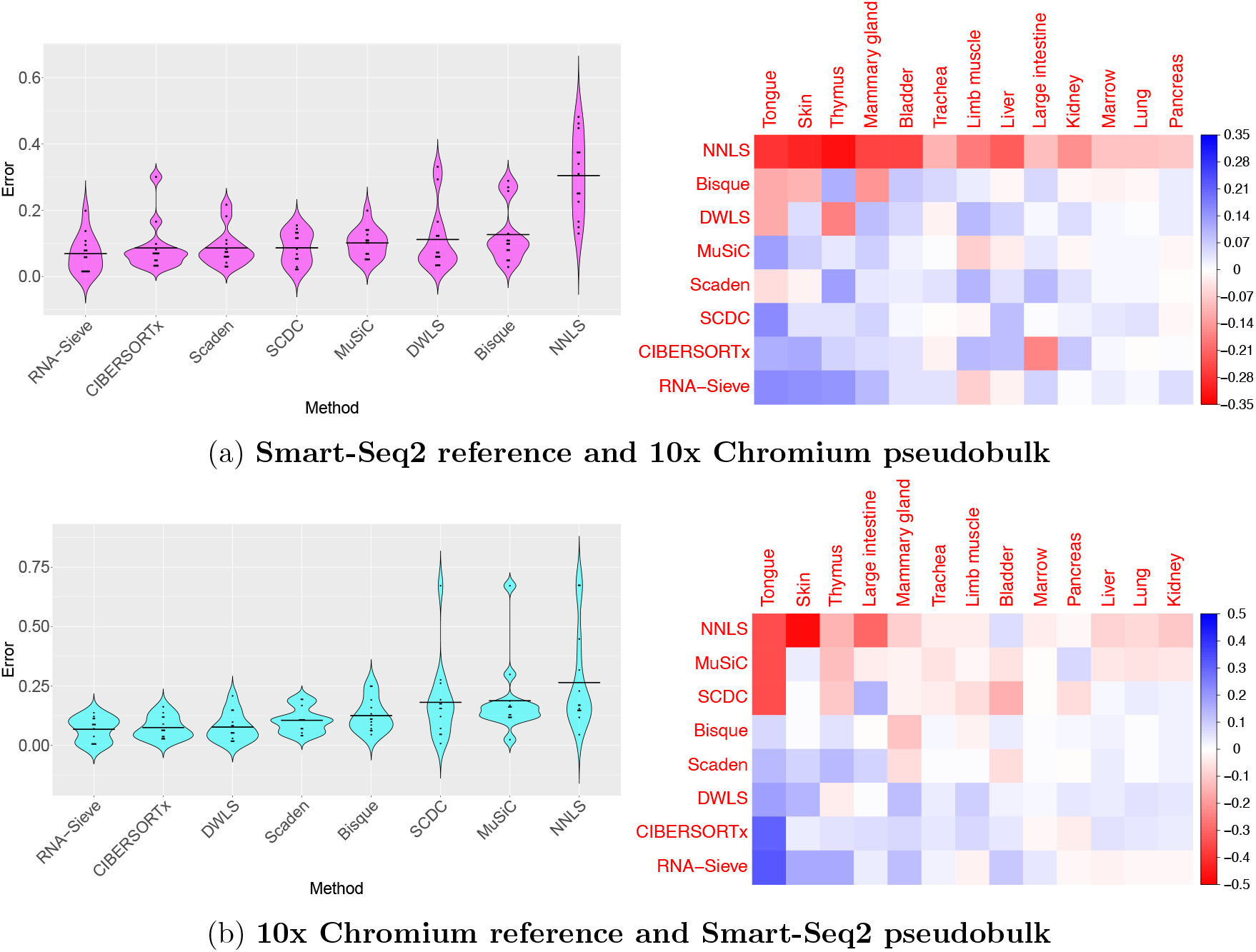
Distribution of errors for each method in pseudobulk experiments. Pseudobulk experiments were performed in 13 different organs using data from the *Tabula Muris Senis* experiment. Errors were computed as the average *L*_1_ error across cell types in each organ. In the violin plots, horizontal black bars correspond to the mean error and methods are ordered left to right from lowest to greatest mean error. In the grid plots, methods and organs were ordered using SVD-induced clustering. Roughly speaking, the methods from top to bottom are characterized by improving performance while the organs from left to right are characterized by decreasing variability in different methods’ performances. Color indicates the difference between the average error across methods in that organ; deeper shades of red (blue) indicate poor (good) relative performance. See Table S2 for context regarding the cell types present in each organ.

Although RNA-Sieve’s nominal improvement in the average per-cell-type *L*_1_ metric may appear minor at first glance, we note that a typical tissue consists of several cell types, and thus the overall error may accumulate rapidly. The constraint that mixture proportions sum to 1 means such reductions in error are likely to be meaningful; when errors in inference are of the same order as the proportion of common cell types, it becomes very easy to arrive at incorrect biological conclusions, especially in more complex tissues with many cell types. We show in Figure S3 a representative example of seemingly minor average *per-cell-type* improvement resulting in comparatively major *individual-cell-type* differences. Other error metrics are more sensitive to different aspects of performance and may detect such improvements more reliably, and are touched upon below. By virtue of having to consider multiple distinct algorithms, tissues, cell types, and experimental protocols, any benchmarking evaluation must necessarily consist of a large number of combinations. Between these many factors and the random nature of the data, it is (even theoretically) nearly impossible for any one algorithm to dominate the others in all situations (as is recognized and discussed in Menden et al. (2020)). We believe that evaluation should therefore focus on aggregate measures of accuracy across many situations. We hence supplement Table 1 and Figure 2 with Table 2, which presents the mean ranks of all eight algorithms aggregated across all (26) cross-protocol experiments using the *L*_1_, *L*_2_, *L*_∞_, and *KL* error quantifications as these accuracy metrics assess different aspects of model performance (see Benchmarking procedures). We find RNA-Sieve outperforms its nearest competitor by roughly one-half rank regardless of error metric, a gap which is at least as large as those between other neighboring methods (NNLS excluded).

**Table 2:**
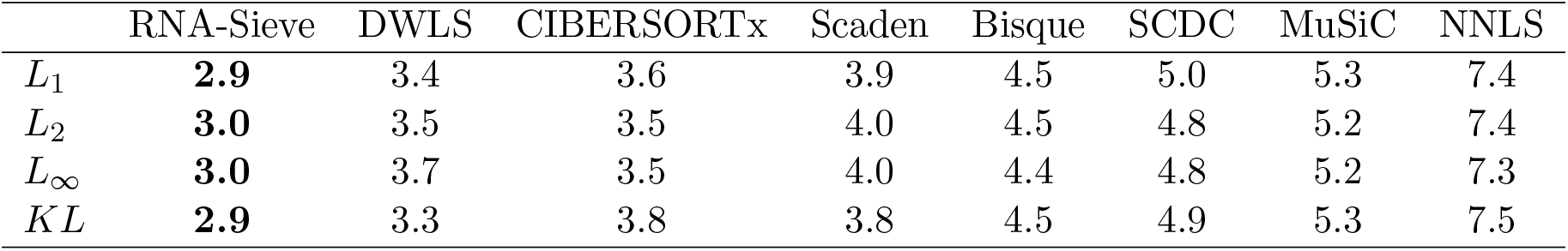
Mean ranking of algorithms under various error metrics. All eight methods were ranked 1 (best) to 8 (worst) on all 26 cross-protocol deconvolutions using the *L*_1_*, L*_2_*, L*_∞_ and *KL* (KL-divergence) metrics, and their mean ranks computed.

|  | RNA-Sieve | DWLS | CIBERSORTx | Scaden | Bisque | SCDC | MuSiC | NNLS |
| --- | --- | --- | --- | --- | --- | --- | --- | --- |
| $L_1$ | **2.9** | 3.4 | 3.6 | 3.9 | 4.5 | 5.0 | 5.3 | 7.4 |
| $L_2$ | **3.0** | 3.5 | 3.5 | 4.0 | 4.5 | 4.8 | 5.2 | 7.4 |
| $L_\infty$ | **3.0** | 3.7 | 3.5 | 4.0 | 4.4 | 4.8 | 5.2 | 7.3 |
| $KL$ | **2.9** | 3.3 | 3.8 | 3.8 | 4.5 | 4.9 | 5.3 | 7.5 |

In practice, complications to the generic deconvolution problem may arise. For example, the scRNA-seq reference data may lack one or more cell types found in the bulk sample, or may even contain extra ones. Such problems are more likely to occur when performing cross-experimental or cross-subject deconvolutions, as we typically must. It is thus important to examine how algorithms perform in these situations. We further recognize the necessity to demonstrate robustness to misspecification for a model-based approach like RNA-Sieve. To do so, we selected the kidney, limb muscle, liver, and marrow due to their representative ranges of cell type number and dissimilarities, and considered all possible configurations containing one extra or missing cell type in the single-cell references. When the reference contains too many cell types, deconvolution schemes should infer proportions near zero for these extra cell types. We found that to be the case with RNA-Sieve (Figures 3 and S4) as long as the extra reference cell type is sufficiently distinct from the other cell types present in the reference. When cell types are highly similar, inferred proportions may be shared among them and might not change substantially upon removal of one of these cell types from the bulk. Meanwhile, when a cell type present in the bulk is absent from the reference, the more likely of these two scenarios, the deconvolution problem becomes overdetermined. Ideally, deconvolution algorithms would move the weight of the removed cell type to those most similar to it. Our empirical results (Figures 4 and S5) indicate that RNA-Sieve tends to do precisely this. In some cases, this means mass transfer to one single cell type, while in others the weight is shared among multiple. This result suggests that in the case of misspecification, RNA-Sieve will still achieve sensible solutions as long as sufficiently representative cell types are captured in the reference set. We note that given the generative nature of our model, a hypothesis test to detect missing cell types is, as opposed to existing methods, within the capabilities of our framework (see Discussion).

**Figure 3:**
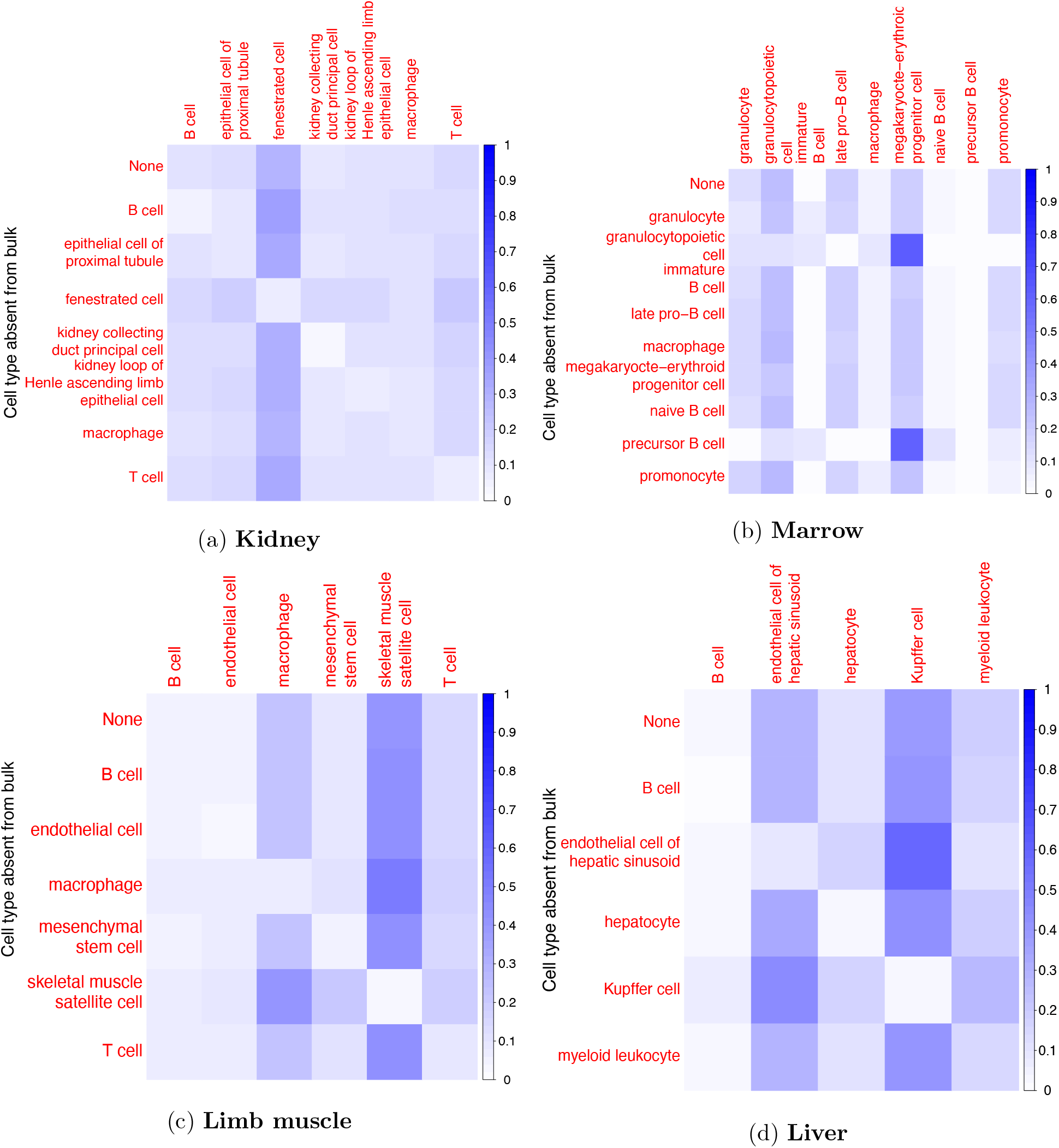
Deconvolution with extra cell types in the reference matrix. Deconvolution was performed in pseudobulk experiments in four different organs from the *Tabula Muris Senis*. For each organ, we followed a leave-one-out procedure in which one cell type is removed from the pseudobulk at a time. Deconvolution was then performed with this extra cell type in the reference in order to examine RNA-Sieve’s specificity. The top row shows the inferred proportions with no extra reference cell types. Darker colors indicate a higher estimated proportion value. Here we used Smart-Seq2 data for the references and 10x Chromium for the pseudobulks.

**Figure 4:**
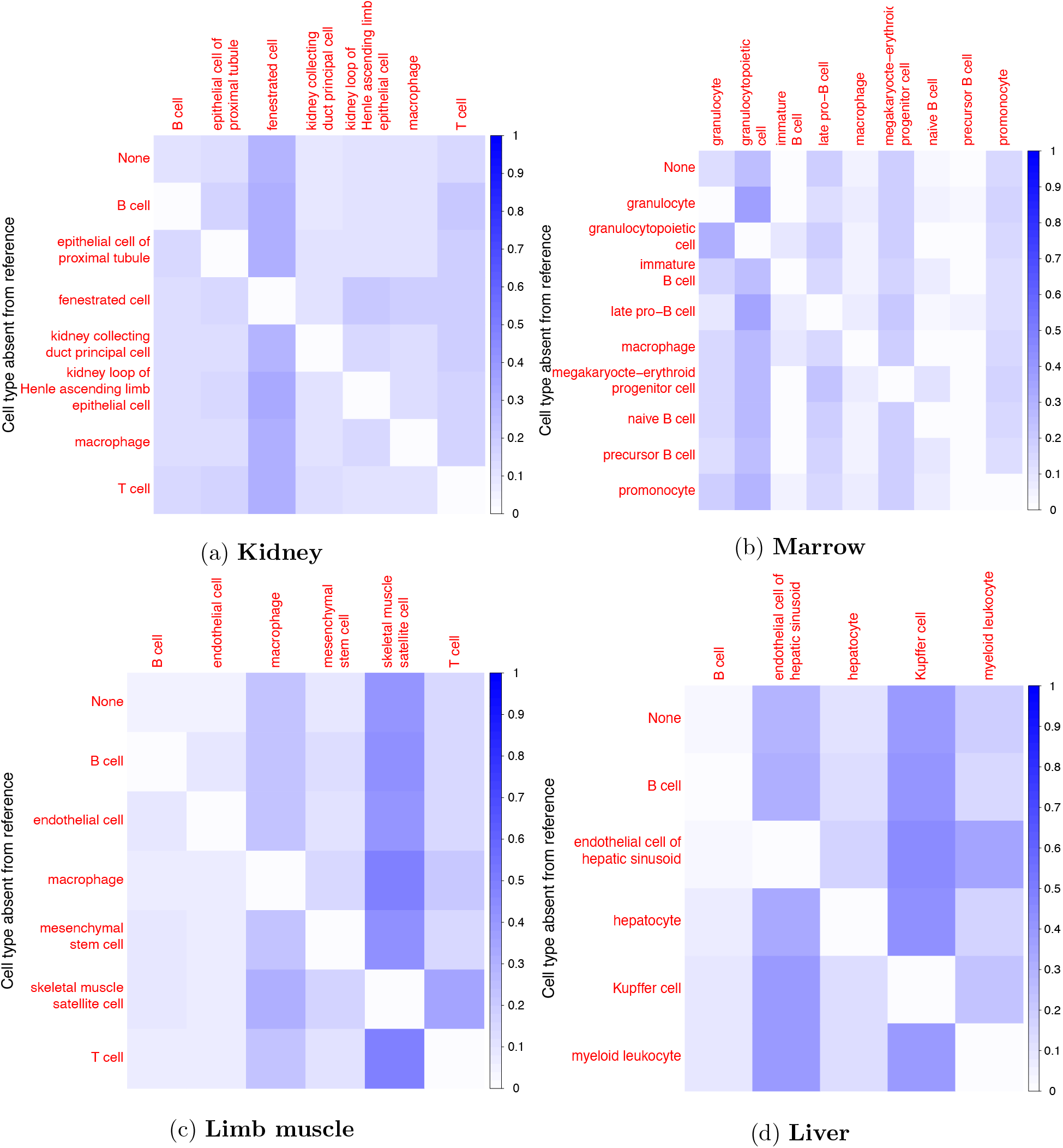
Deconvolution with missing cell types in the reference matrix. Deconvolution was performed in pseudobulk experiments in four different organs from the *Tabula Muris Senis*. For each organ, we followed a leave-one-out procedure in which one cell type is removed from the reference at a time. Deconvolution was then performed with an extra cell type in the pseudobulk in order to examine RNA-Sieve’s ability to handle such a misspecification. The top row shows the inferred proportions with no missing reference cell types. Darker colors indicate a higher estimated proportion value. Here we used Smart-Seq2 data for the reference and 10x Chromium for the pseudobulk.

### Validation with Real Bulk RNA-seq Data

In certain rare instances, bulk RNA-seq data sets with known or experimentally estimated cell type proportions are available. We considered three such data sets in order to evaluate RNA-Sieve under more realistic conditions. The first of these data sets is a bulk RNA-seq mixture of two human breast cancer cell lines and fibroblasts with accompanying scRNA-seq data first published in Dong et al. (2020). These cells were mixed together in proportions of 60% MDA-MB-468, 30% MCF-7, and 10% fibroblasts. As shown in Table 3, RNA-Sieve yields highly accurate results, attaining the lowest average error among all methods. With the exception of SCDC, other methods overestimated the fraction of the MCF-7 cell line in the bulk while underestimating the MDA-MB-468 cell line by large amounts, and most methods substantially overestimated the fibroblast proportions.

**Table 3:** Inferred proportions from different methods in cell line mixture experiment. Data from Dong et al. (2020) with known cell type proportions was used to evaluate each applicable method (displayed proportions may not sum precisely to 1 due to rounding). Bisque and MuSiC are not intended for use with only one individual in the bulk data and/or single-cell reference and were thus not included. SCDC was run in tree mode for this deconvolution.

| Method | Estimated Proportions (%) | | | Mean $L_1$ Error |
| --- | --- | --- | --- | --- |
|  | 60% MDA-MB-468 | 30% MCF-7 | 10% Fibroblasts |  |
| RNA-Sieve | 62 | 26 | 13 | 3 |
| SCDC | 60 | 19 | 21 | 4 |
| Scaden | 35 | 44 | 21 | 17 |
| CIBERSORTx | 32 | 52 | 16 | 19 |
| DWLS | 26 | 48 | 27 | 23 |
| NNLS | 22 | 56 | 21 | 25 |

Because this experiment contains only three cell types and a single bulk sample all from one experiment, we sought to validate using larger data sets containing expression measurements from peripheral blood mononuclear cells (PBMCs) which promised to be more heterogeneous. The first of these, analyzed in Newman et al. (2019), measures gene expression in twelve bulk samples and a scRNA-seq reference from one individual. Ground-truth cell type proportions in all bulk samples were estimated using flow cytometry and were grouped into six primary categories: B cells, CD4+ T cells, CD8+ T cells, monocytes, natural killer (NK) cells, and neutrophils. The second PBMC bulk data set comes from Monaco et al. (2019) and contains a further twelve individuals, with flow cytometry again providing cell type proportion estimates for the same cell types. We obtained two scRNA-seq PBMC reference data sets. The first, which we used with the Newman et al. bulk, also comes from Newman et al. and assays one individual. To explore the effect of multiple individuals in the reference, we downloaded two reference sets from the public repository managed by 10x Genomics and subsequently merged them; this reference was used with the Monaco et al. bulk data samples. As neutrophils are notably difficult to assay accurately at the single-cell level, they were not present in either of the original reference panels. However, given the large fractions of neutrophils estimated by flow cytometry, particularly for the Newman et al. data set, we identified a publicly available data set which contains scRNA-seq data for human neutrophils (Xie et al., 2020). These data were then incorporated into both reference sets in order to enable more effective comparisons. Because the Newman et al. scRNA-seq reference was relatively small (tens to hundreds of cells per cell type) and only had one individual present, we subsampled the neutrophil data down to 250 cells from one individual to be consistent with the other cell types. Conversely, because the 10x Genomics PBMC referece had more cells (hundreds to thousands of cells per cell type) and multiple individuals, we subsampled 1250 neutrophils in total from three individuals for use in the reference (see Benchmarking procedures).

Subsequent deconvolutions showed that RNA-Sieve performed the best out of all methods as measured by the mean absolute deviation (*L*_1_ error) when aggregating across both analyses (Table 4). The results are summarized graphically in Figure 5. The presence of neutrophils presented a challenge for several methods, perhaps due to the fact that they came from a different experiment or because of their uniquely low RNA counts. For example, in the bulk data from Newman et al., neutrophils were strongly underestimated by CIBERSORTx, Scaden, and SCDC with most of that mass being allocated to either monocytes, CD4+ T cells, or B cells, respectively. RNA-Sieve and DWLS both performed well on these bulk samples, though RNA-Sieve slightly underestimated neutrophils in favor of monocytes while DWLS had minor difficulty distinguishing between CD4+ and CD8+ T cells. A similar story emerged for the Monaco et al. data, with CIBERSORTx, DWLS, Scaden, and, to a lesser extent, RNA-Sieve underestimating neutrophil and CD8+ T cell proportions while overweighting monocytes or CD4+ T cells. In contrast, Bisque, SCDC, and MuSiC strongly overweighted neutrophils (and sometimes natural killer cells) at the expense of other cell types. To produce a more formal and comprehensive comparison, we computed summary statistics in the same manner as Table 2 using the 24 bulk samples comprising the two data sets. RNA-Sieve achieves the best performance among all considered methods (Table 5) in each metric as it exhibits strong performance for both data sources. DWLS performs well on the Newman et al. data but fails to attain that level of accuracy on the Monaco et al. data.

**Table 4:** Average *L*_1_ errors with PBMC data and ground-truth cell proportions from flow cytometry. The first two columns display average *L*_1_ errors for the two PBMC data sets individually, while the last column aggregates *L*_1_ errors across both data sets. Bisque and MuSiC do not provide proportion estimates for the Newman et al. data because only one individual is present for all reference cell types. CIBERSORTx was run in B-mode per their recommendation with a UMI-based scRNA-seq reference.

| Method | Average $L_1$ Error (%) | | |
| --- | --- | --- | --- |
|  | Aggregate | Newman et al. Data | Monaco et al. Data |
| RNA-Sieve | **4.8** | 4.8 | **4.7** |
| DWLS | 7.2 | **4.7** | 9.7 |
| Scaden | 9.4 | 11.3 | 7.6 |
| CIBERSORTx | 14.4 | 17.7 | 11.2 |
| SCDC | 19.3 | 17.2 | 21.3 |
| NNLS | 25.2 | 27.7 | 22.7 |
| Bisque | n/a | n/a | 17.3 |
| MuSiC | n/a | n/a | 22.7 |

**Table 5:**
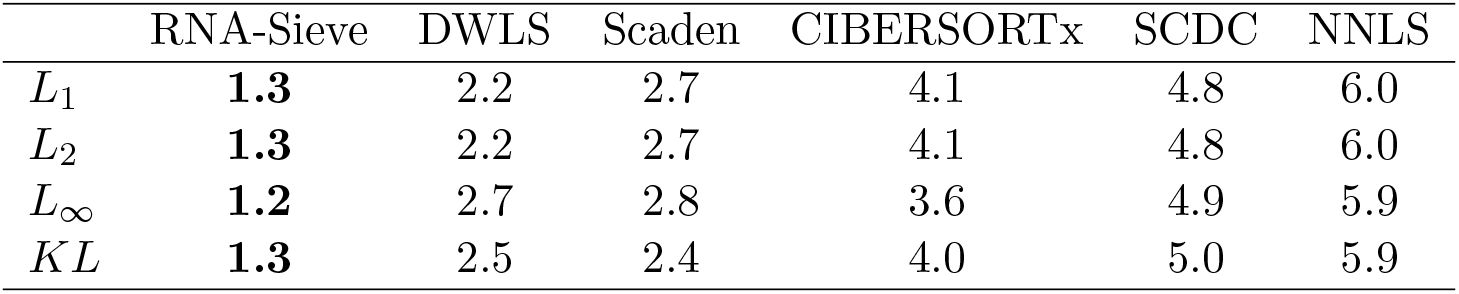
Mean ranking of algorithms under various error metrics combined across the two PBMC deconvolutions. All six applicable methods were ranked 1 (best) to 6 (worst) across 24 bulk samples from the Newman et al. and Monaco et al. data using the *L*_1_*, L*_2_*, L*_∞_ and *KL* (KL-divergence) metrics, and their mean ranks were computed.

|  | RNA-Sieve | DWLS | Scaden | CIBERSORTx | SCDC | NNLS |
| --- | --- | --- | --- | --- | --- | --- |
| $L_1$ | **1.3** | 2.2 | 2.7 | 4.1 | 4.8 | 6.0 |
| $L_2$ | **1.3** | 2.2 | 2.7 | 4.1 | 4.8 | 6.0 |
| $L_\infty$ | **1.2** | 2.7 | 2.8 | 3.6 | 4.9 | 5.9 |
| $KL$ | **1.3** | 2.5 | 2.4 | 4.0 | 5.0 | 5.9 |

**Figure 5:**
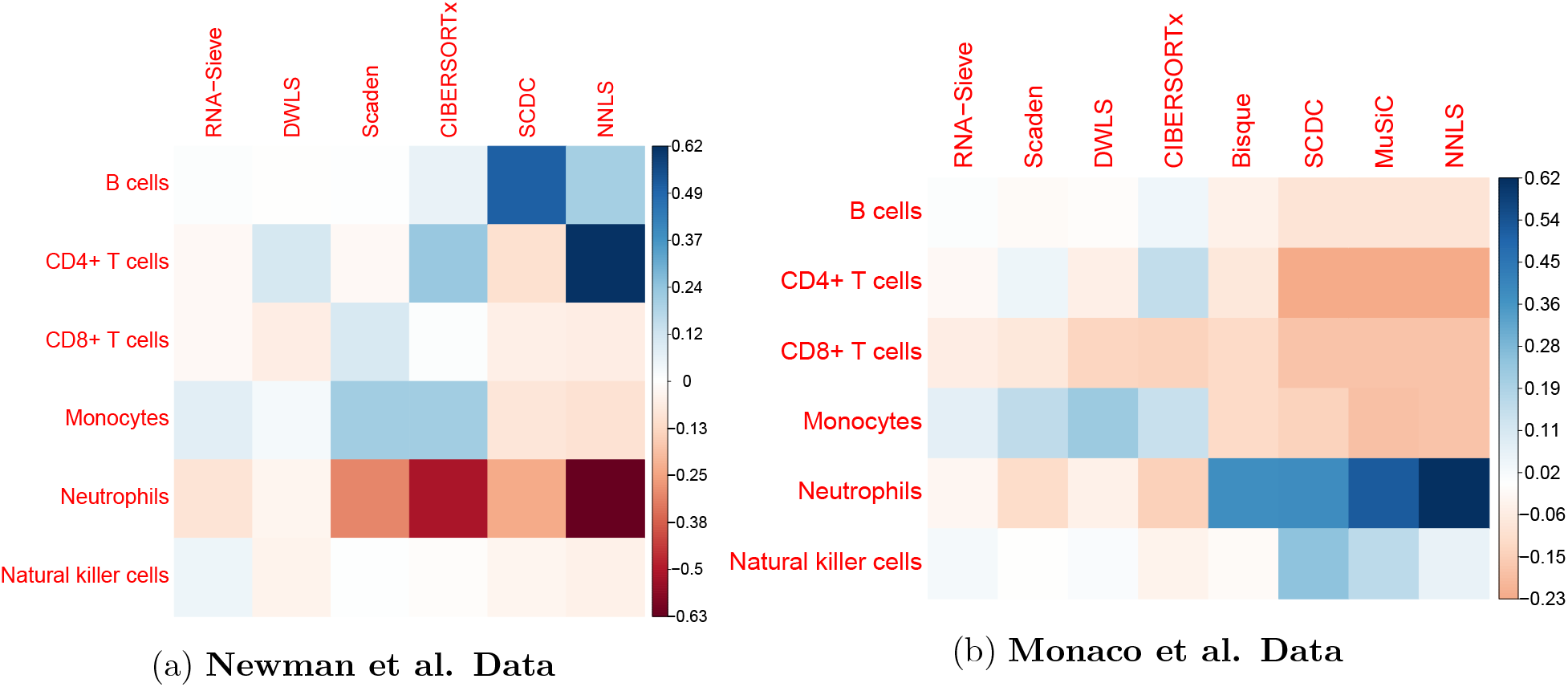
Deconvolution biases for PBMC data with known ground-truth proportions. Average differences between inferred and true proportions were computed within each cell type across the twelve bulk samples present in each scenario. Consistent overestimation of a cell type’s abundance results in darker blue squares, while red corresponds to chronic underestimation. Methods are ordered left-to-right by overall performance.

Finally, we analyzed samples from the pancreatic islets region of the human pancreas where ground-truth proportions were not available. This region has previously been used for validation in the absence of ground-truth proportions because of prior knowledge of the general ranges of constituent cell types. Moreover, the well-known negative relationship between beta cell proportions and hemoglobin A1c (Hb1Ac) levels allows us to test whether different deconvolution approaches can recapitulate this relationship. As shown in Figure S6, RNA-Sieve is among the methods which successfully identify the expected negative correlation. As ground-truth values were not available for these data, it is impossible to ascertain precisely how methods performed, though it appears each method’s average inferred beta cell proportions are below the expected ~ 50%. Nevertheless, successful recovery of the expected association between beta cell proportions and Hb1Ac levels serves as a useful benchmark. Given the necessity to demonstrate robust performance across a range of tissues and cell type groups, we feel this result provides important support to RNA-Sieve’s strong performance in the cell line and PBMC deconvolution tasks above.

### Analysis of Real Bulk Organ Samples

We next applied RNA-Sieve to real bulk RNA-seq data to look for interesting patterns in organ composition. We chose to continue working with the *Tabula Muris Senis* data set as it contains many bulk RNA-seq samples in addition to the scRNA-seq data previously described. Due to its expansive experimental design across organs and ages, this resource is uniquely suited to interrogate changes in cell type compositions associated with the process of aging. By identifying changes in the balance of cell classes, we hope to provide insight into shifts in the mechanisms driving organ functions at different stages of life. In general, aging represents one of the more complicated biological processes, and one which occurs in every person or organism. Due to its ubiquity and significant effects on quality of life, improved understanding of the etiologies underlying age-associated functional deficits holds great potential therapeutic value. Degradation of the musculoskeletal and immune systems are among the most apparent phenotypic changes occurring during mammalian aging. Here we highlight intriguing results from three organs with roles in these bodily systems–the limb muscle in the former, and the spleen and bone marrow in the latter. In the absence of ground truth, we judge the reliability of our estimates by the relative consistency of inferred proportions within and across age groups.

Limb muscle in the arms and legs provides support and locomotion. It primarily consists of skeletal muscle, stromal, immune, and endothelial cells. Differentiated muscle cells contract to produce the aforementioned support and motion, while satellite cells serve in myogenesis and muscular repair (Christov et al., 2007). Stromal cells comprise the connective organ which binds sarcomeres together and connects muscles to bones in addition to displaying certain regenerative capabilities (Kasprzycka et al., 2019). Immune response is often noted in muscle cells after mechanical stress-induced damage to muscle fibers, when the tissue becomes inflamed due to tearing (Novak et al., 2014; Burzyn et al., 2013). Endothelial cells are present due to the often high degree of vascularization needed to support muscle function (Christov et al., 2007). Upon application of RNA-Sieve to the available bulk muscle samples, we observed a noticeable increase in skeletal muscle satellite cells and a substantial decrease in the mesenchymal stem cell proportion in older mice (Figure 6a). These trends are present, albeit fairly gradual, until around 21 months old with more sudden changes apparent thereafter. There was also an apparent increase in macrophage proportions up until 15 months of age, followed by a slow decline for the remainder of life. Each of these three cell types has been demonstrated to function in muscle fiber repair through different mechanisms (Snijders et al., 2015). This pattern in cell type composition may thus indicate changes in the relative use of different regenerative pathways as individuals age.

**Figure 6:**
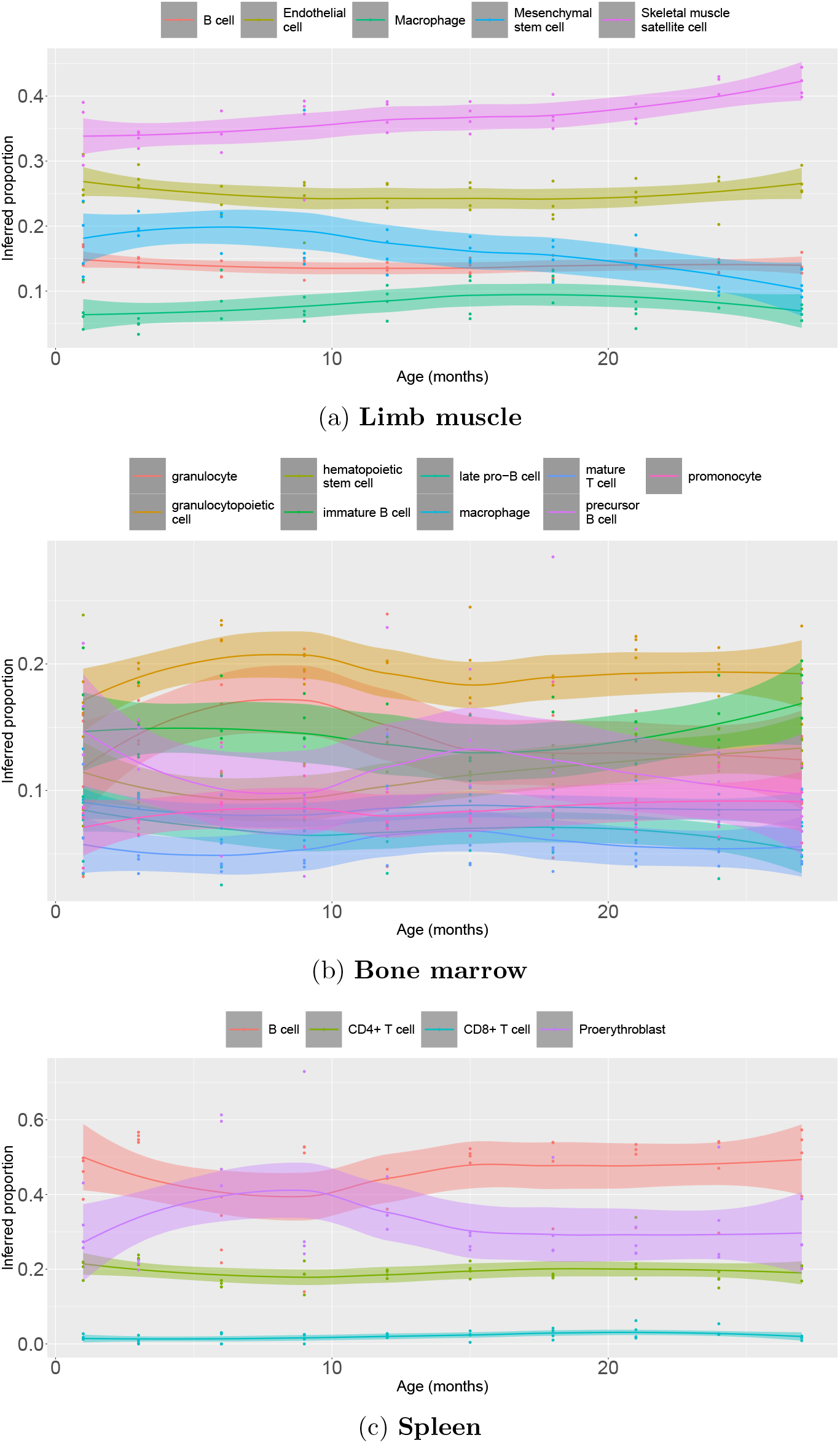
Deconvolution results for real bulks from the *Tabula Muris Senis*. Roughly forty real bulk samples across ten ages were deconvolved using RNA-Sieve in each of the limb muscle, bone marrow, and spleen. In all cases, Smart-Seq2 data were used as the reference. Each point represents the inferred proportion for a given cell type in a bulk sample. Lines display the smoothed trend of proportions as a function of age, with uncertainty shown by the shaded intervals.

The bone marrow is a vital component of the immune system which executes the bulk of hematopoiesis and contains numerous constituent cell types ranging from various stem cells to more mature cell classes (Gurkan and Akkus, 2008). This rich combination yielded several age-associated trends in cell type composition, and we choose to focus on two. First, an effectively linear growth in the number of hematopoietic stem cells was observed with increasing age. Though this may seem surprising given reduced adaptive immunity with age, this exact phenomenon has been previously observed in both mice and humans (Pang et al., 2011), and it is accompanied by a decrease in functionality of these cells. Conversely, granulocyte proportion appeared to decrease after roughly 9 months of age. Further examination reveals that the granulocyte fraction tends to mirror that of granulocytopoietic cells, but with an increasing deficit between the two as age increases. Such a pattern is suggestive of the reduced potency of granulocytopoiesis that we would expect with age. Hence, in the marrow we are able to identify known patterns of cell type composition variation despite the presence of many transcriptomically similar cell types.

The spleen occupies a central role in the lymphatic system and is important, though not strictly essential, for proper functioning of the immune system and red blood cell recycling. This organ is split into two pieces—the red pulp, which contains blood cells, and the white pulp, which is primarily lymphatic (Mebius and Kraal, 2005). Typically, the large majority of present immune cells are B and T cells, with smaller quantities of other cell types (Hensel et al., 2019). Various progenitor cells may be present to spur production of immune and red blood cells, though these processes are primarily performed in the bone marrow (Mumau et al., 2018). It is notably difficult to distinguish among these progenitor cells in their early stages, making it possible that several varieties are captured within the single label of proerythroblasts. Upon deconvolution, we found that our inferred proportions for B and T cells matched accepted ranges (Hensel et al., 2019). More interestingly, we noticed an unexpected and transient spike in the proportion of proerythroblasts peaking at roughly nine months of age (Figure 6c). Importantly, this increase is observed in all four of the 9-month-old individuals, and is thus not an artifact of outlying samples. Mice at this age are roughly analogous to humans of between 30-40 years of age, and as hematopoiesis is generally restricted to the marrow at this age except under stress conditions, it is unclear whether a programmed hematopoeitic process is occurring or if we are capturing the behavior of a cell type not enumerated in the reference set.

### Extension to Confidence Regions

Within the deconvolution task, the generative framework of RNA-Sieve permits extensions which remain out of reach using prior approaches. One such possibility is the computation of confidence regions for inferred cell type proportions. Despite its clear importance, error quantification in deconvolution is challenging and has received relatively scant attention, leaving users to only guess at the reliability of their results. As deconvolution is sometimes performed upstream of tasks such as differential expression or eQTL detection, it is critical to know whether inferred proportions are precise. Because RNA-Sieve infers these proportions via maximum likelihood estimation, we can directly tap into the wide array of theory on asymptotic confidence bounds. Specifically, we construct confidence regions for inferred proportion values through numerical computation of the inverse empirical Fisher information matrix (see Confidence Intervals in Methods). We demonstrate RNA-Sieve’s ability to produce well-calibrated confidence regions by constructing them for within- and cross-protocol pseudobulk deconvolutions using *Tabula Muris Senis* data as well as both real PBMC bulk data sets in Validation with Real Bulk RNA-seq Data.

We began with within-protocol comparisons where all modeling assumptions are generically met. As shown in Figures 7a and S7a, we obtain narrow, yet well-calibrated, confidence intervals, indicating the effectiveness of our procedure in this simplified scenario. However, the typical deconvolution setting will present complications in the form of protocol differences in the scRNA-seq reference and bulk RNA-seq data. Under mild and plausible assumptions on these distributional shifts, our MLE framework is robust to such model misspecification (see Confidence Intervals), and we still achieve good performance in spite of protocol mismatch (Figures 7b and S7b). Aggregating across runs, our 95%-confidence intervals contain the true cell type proportions 96.7% and 91.8% of the time in the within- and across-protocol deconvolutions, respectively.

**Figure 7:**
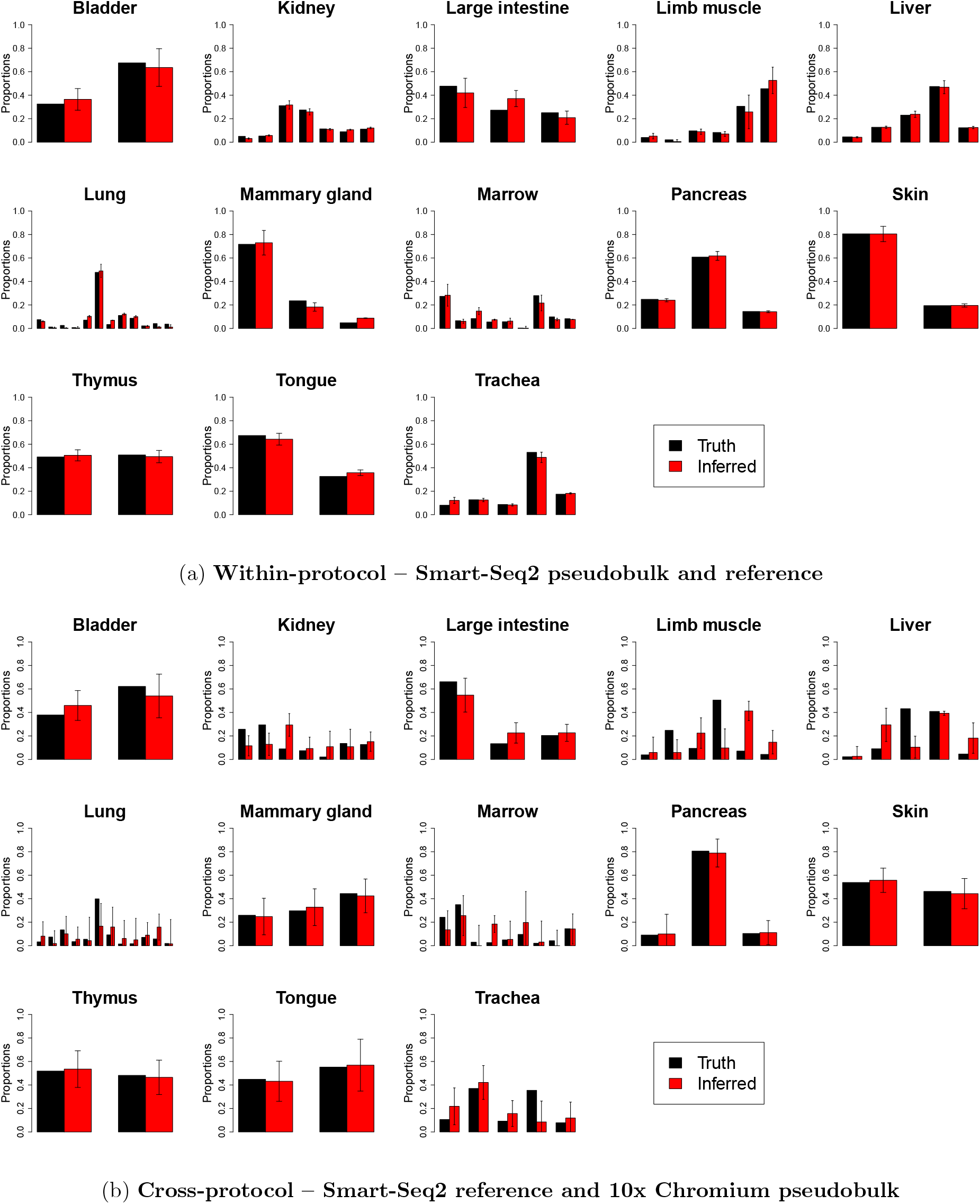
RNA-Sieve results with confidence intervals in pseudobulk experiments. Inferred cell type proportions in pseudobulk experiments using data from the *Tabula Muris Senis* experiment. The black error bars on inferred proportions show the marginal 95% confidence intervals as computed from the estimated Fisher information produced by RNA-Sieve. See Confidence Intervals for mathematical details. Table S2 contains the cell types in each organ, which could not be displayed because of space constraints.

To ensure that we obtain sensible results with real bulk RNA-seq data, we also generated confidence intervals for the whole blood samples analyzed in Validation with Real Bulk RNA-seq Data. We again obtain calibrated and sensible results, with our confidence intervals containing the truth 95.8% of the time in the Monaco et al. bulk samples and 90.3% of the time in the Newman et al bulk samples (Figure 8). Though assessing their accuracy is impossible absent ground-truth proportions, we also computed confidence intervals for the real bulks deconvolved in Analysis of Real Bulk Organ Samples to verify that RNA-Sieve’s confidence intervals were reasonable in tissues besides whole blood. We found that these interval widths were similar to those we obtained in our other trials (Figure S8). The distribution of confidence interval half-widths for cell type proportions were also generally consistent across samples (Figure S9). We note that MuSiC presents a quantity which seemingly corresponds to the variance in proportion estimates, though it was not emphasized in their manuscript, and we generally found the produced values to be overly small in practice.

**Figure 8:**
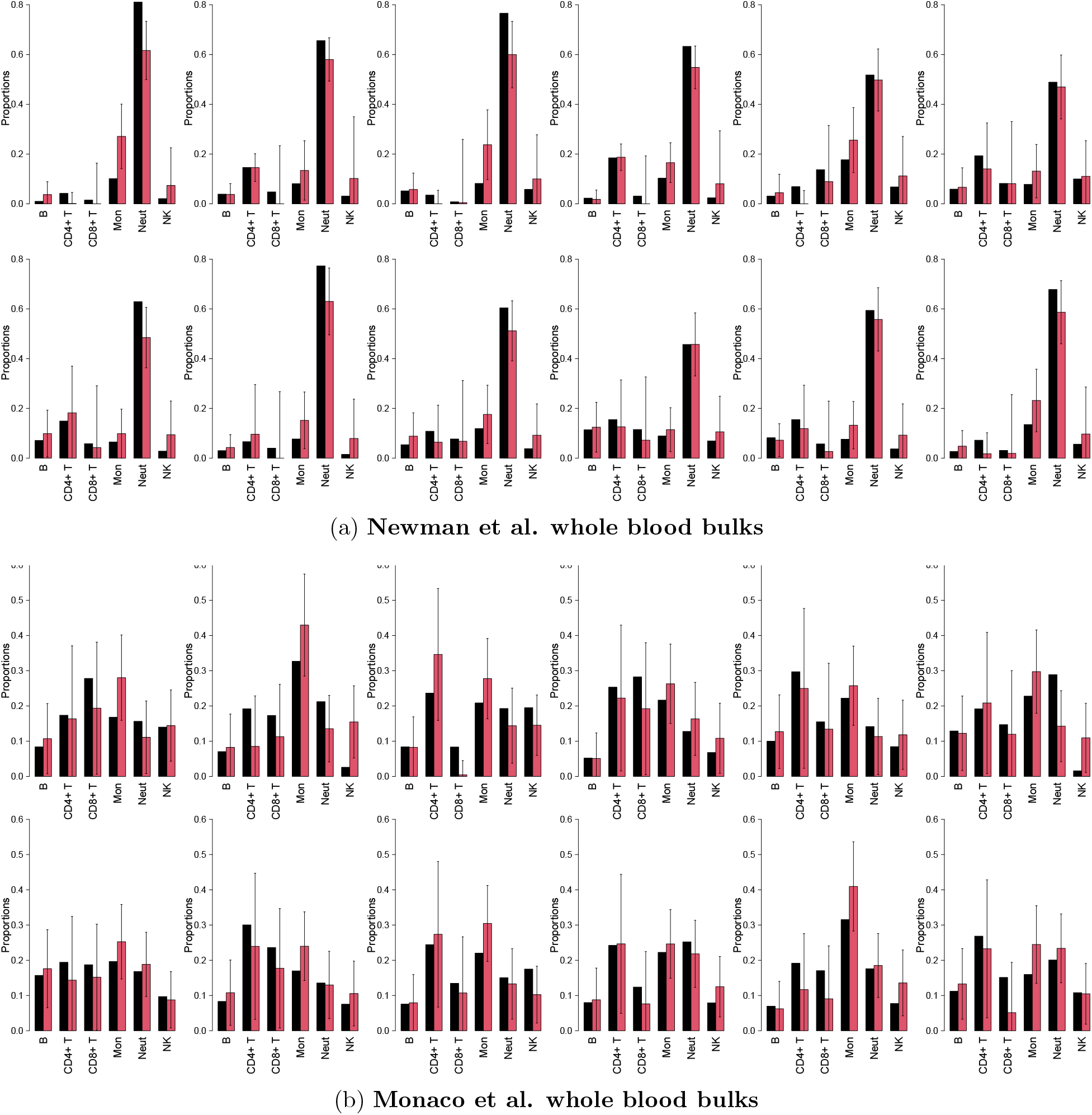
RNA-Sieve results with confidence intervals for whole blood bulk samples with known cell type proportions. Inferred cell type proportions in deconvolutions using PBMC references and whole blood bulks as described in Validation with Real Bulk RNA-seq Data. True proportions as estimated by flow cytometry are in black while RNA-Sieve’s inferred proportions are in red. The black error bars on inferred proportions show the marginal 95% confidence intervals as produced by RNA-Sieve. Confidence intervals capture true proportions 90.3% and 95.8% of the time in the respective scenarios. See Confidence Intervals for mathematical details.

In principle, the widths of confidence intervals should depend on the number of cells and genes in the reference, similarity among cell types in the reference, and agreement between the reference and bulk measurements. Our empirical results suggest that these eminently logical factors do, in fact, drive the widths of our intervals. For example, the confidence intervals in cross-protocol deconvolutions are a bit wider than their within-protocol counterparts, due to our adaptive procedure’s conservative nature when it detects differences between the reference and bulk. This arises in part because we deem fewer genes reliable when compared to within-protocol experiments. Moreover, evidence of the contribution of reference sample size is present in a few organs, most notably the lung with its many low-frequency cell types.

## Discussion

Here we have introduced our method for supervised bulk gene expression deconvolution, RNA-Sieve, and illustrated its robust performance in a variety of settings. Unlike methods which rely on variants of least squares or the application of complex machine learning algorithms, we place the deconvolution problem into a generative probabilistic framework that models random noise in both the reference panel and bulk samples by relying on asymptotic theory. Through simulations and applications to real data, we demonstrated the broad applicability of our method and its utility to investigate biological questions of interest.

It is valuable to understand how RNA-Sieve differs from other approaches and to consider the consequences of these divergent design choices. Least squares-based solutions such as MuSiC (Wang et al., 2019), SCDC (Dong et al., 2020), and DWLS (Tsoucas et al., 2019) devise their own implementations of weighted non-negative least squares (W-NNLS). These methods aim to handle heteroskedasticity across genes by re-weighting them according to their variability and specificity, allowing genes which are putatively more informative to carry increased importance in the regression task. Alternatively, Bisque (Jew et al., 2020) uses NNLS after applying a transformation to bring the reference and bulk data into better distributional agreement. From a modeling perspective, least squares-based solutions generally address uncertainty in the bulk only, leaving stochasticity in the single-cell reference unaccounted for. With RNA-Sieve, rather than devising a specialized gene-weighting scheme, we naturally emphasize some genes more than others via variances resulting from an explicit generative model incorporating noise in both single cells and the bulk. We also do not explicitly attempt to bring reference and bulk data into better agreement a là Bisque, instead relying on a filtering protocol to remove genes which display signs of significant deviation from our assumptions. Integrating an explicit transformation is an interesting possibility for RNA-Sieve, and should only boost its performance by further aligning data to our model assumptions. Other methods employ machine learning techniques, such as CIBERSORTx (Newman et al., 2019), which uses *ν*-support vector regression, and Scaden (Menden et al., 2020), which utilizes deep neural networks. These approaches can be opaque to the user due to their reliance on high-complexity algorithms which often lack theoretical guarantees of optimality and provably accurate inference despite continuing advances in explainability techniques. Comparatively, our formulation of RNA-Sieve as the MLE of an explicit generative model is transparent in both parameter interpretation and performance guarantees. The parameters updated during optimization have explicit biological meanings and tracing their values allows for a deeper interrogation of the predictions RNA-Sieve generates. This is a useful feature when providing context to inferred cell type proportions as well as exploring the theoretical limits of deconvolution as a function of cell type properties. Like MuSiC, SCDC, Bisque, and Scaden, we do not select marker genes in RNA-Sieve. This helps us maintain computational efficiency, while simultaneously providing robustness with respect to outlier fluctuations in gene expression. We also parallelize our optimization steps and jointly update parameters when deconvolving multiple bulk samples. This yields significant speedups relative to serial runs and allows us to share statistical strength across all bulks.

RNA-Sieve is embedded in a flexible generative framework, which can be adapted to a variety of situations to make deconvolution performance more effective. One of these is the modeling of further sources of variation. For instance, if gene expression distributions are expected to differ drastically across individuals from which samples are taken, this knowledge can be explicitly incorporated into our likelihood. Without such modification, RNA-Sieve implicitly follows the paradigm of MuSiC, SCDC, and Bisque in penalizing genes of large inter-individual variance via the marginal variances resulting from estimation of the reference panel. A similar notion applies to mitigating potential batch effects or effectively combining disparate references. Currently, different reference matrices which are believed to have the same expression distributions can be averaged together to increase statistical power without further modification of our present implementation.

A principal motivation of this work was to expand the scope of accessible questions in the deconvolution setting. Our likelihood-based approach facilitates extensions which are intractable with current algorithms. As a first step, we have chosen to demonstrate our ability to explicitly construct confidence regions for inferred proportions, producing a mathematically rigorous quantification of the uncertainty in our estimates. The necessity of these bounds is plainly substantiated by the use of deconvolution upstream of tasks ranging from cell type-specific differential expression to eQTL detection using heterogeneous RNA-seq organ samples. The credibility of any such analyses is predicated on the accuracy of deconvolution, because any errors in this initial step will propagate through to the final result. Consequently, we anticipate that our confidence regions will encourage improved assessment of the reliability of results obtained through these types of analyses. Our confidence intervals are also of obvious inherent value when using deconvolution results to infer differences in cell type composition between samples, whether due to disease status or other factors. Beyond error quantification via confidence intervals, potent possibilities lie in hypothesis testing. Currently, CIBERSORTx does propose one type of test, though our understanding is that it tests whether *any* of the bulk cell types were found in the reference. This is rather restrictive, so we hope to develop procedures with broader utility. One example with clinical impact is a test to determine whether the reference panel is missing cell types present in the bulk sample. Even though we have demonstrated that RNA-Sieve is robust with respect to such misspecification (see Performance in Pseudobulk Experiments), it is nonetheless beneficial to know whether the deconvolution performed was sufficiently valid using a principled approach. Such a test can be directly developed in our framework by examining the residuals produced by our maximum likelihood estimate, and work in this direction is underway.

Despite the flurry of recently developed methods, the question of statistical deconvolution of gene expression data remains far from solved. RNA-Sieve illustrates the efficacy, adaptability as well as promise of generative modeling in this setting, and we hope it spurs continued development within other methodological paradigms. In particular, notions of error quantification and hypothesis testing merit further attention.

## Methods

Below we present the mathematical details that justify our design choices in formulating RNA-Sieve. For the reader interested in more high-level guidance on when to use RNA-Sieve and what potential preprocessing steps to take, we compiled Table S3 as an accessible overview.

### Notation

To ease parsing of technical equations, we briefly introduce our notation here. We generally refer to vector quantities with boldfaced lowercase letters, while plain lower- and uppercase symbols are reserved for scalars (or scalar functions) and matrices, respectively. The *k*^th^ column vector of a matrix 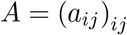 is written as ***a****_k_*, and inner products between vectors ***v***, ***w*** are typically denoted ⟨***v***, ***w***⟩. To distinguish observed, random quantities from their underlying deterministic, ground truth objects we add tildes to the former and asterisks to the latter; e.g., 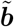 are observed bulk gene expres-sions, while ***b***\* are the true bulk gene expression means. Estimates of latent parameters carry hats; e.g., 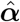 is the vector of mixture weights inferred byour dceonvolution procedure. Finally, we denote by [*n*] the set of *n* elements {1*, …, n*}, and by Δ^*K* − 1^ = {*x* ∈ ℝ^*K*^: ||*x*||_1_ = 1 and *x_k_* ≥ 0 for all *k*} the *K* − 1 dimensional simplex.

### Mathematical Model

We assume that for each gene *g* ∈ [*G*] and cell type *k* ∈ [*K*], there exists a distribution *ν_g,k_* describing the expression of gene *g* in cell type *k*. As multiple cell types comprise any given organ/tissue, the expression of gene *g* in a cell drawn at random from a organ/tissue is governed by the mixture distribution

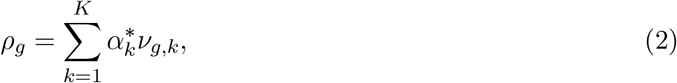

where 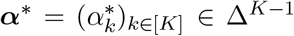 contains the proportions of each cell type in the organ/tissue of interest. Despite the *a priori* infinite-dimensional setting, if *G > K* and *ρ_g_*, 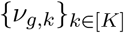 are fully known and sufficiently distinct, the convex combination of (2) immediately implies that ***α***\* can be recovered as the unique solution of the finite-dimensional problem

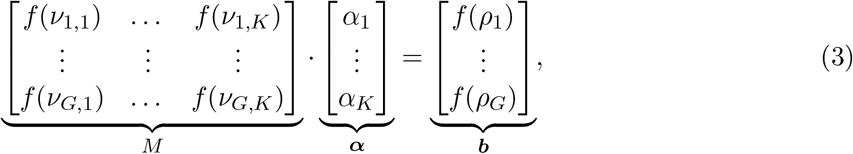

where *f* is any suitable linear function on the space of probability distributions on ℝ (i.e., *f* (Σ_*j*_ *w_j_ μ_j_*) = Σ_*j*_ *w_j_ f* (*μ_j_*) for any convex combination of distributions *μ_j_*). Natural *f* to consider include point evaluations at *x* ∈ ℝ; i.e., *f(ν)* = *F_ν_(x)*, where *F_ν_* denotes the cumulative distribution function (CDF) of *ν* or its *i^th^* moments *f(ν)* = ∫ *x^i^ν(dx)*, both of which enjoy a wealth of statistical theory and proposed estimators.

In experimental settings, exact gene expression distributions are not accessible and instead must be estimated, so utilizing easily and robustly inferrable *f* becomes crucial. In addition to not having direct access to 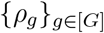, any analysis is further complicated by the fact that bulk sequencing only yields gene expression levels over whole samples and not for particular cells or cell types. That is, the output is effectively a random variable 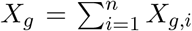 where 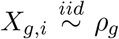 gives the measured expression of gene *g* aggregated over the *n* ∈ ℕ individual cells comprising the sample. It is thus expedient to choose an *f* in (3) that is not only linear on the space of probability distributions, but also for sums of random variables. The essentially unique such *f* is the expectation 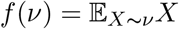, which turns (3) into

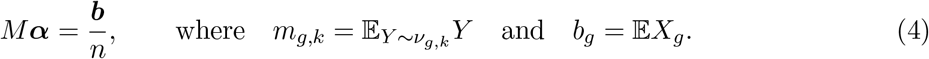

Incorporating the fact that we only observe noisy bulk samples *X_g_* instead of *b_g_* directly results in

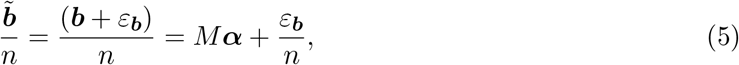

where 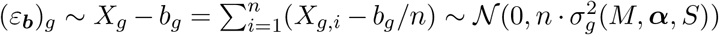 for large *n* by the central limit theorem (CLT), with 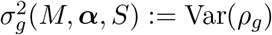 as a function of *M*, ***α***, and 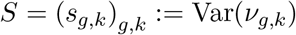.

### Incorporating the dependence of 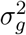 on *α*

If the dependence of 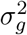 on ***α*** is ignored, (5) lends itself to a simple (weighted) non-negative least squares scheme solving

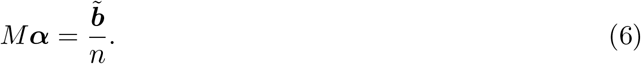

This yields a solution 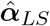 of roughly 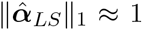 that simply requires re-scaling to fit onto the simplex. This together with data-driven modifications is the approach pursued in Dong et al.(2020); Tsoucas et al. (2019); Wang et al. (2019), where it is argued that (6) outperforms previous methods.

The first improvement of RNA-Sieve over previous approaches stems from explicitly incorpo-rating the dependence of 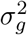 on ***α***. More concretely, we first make explicit this ***α***-dependence by computing

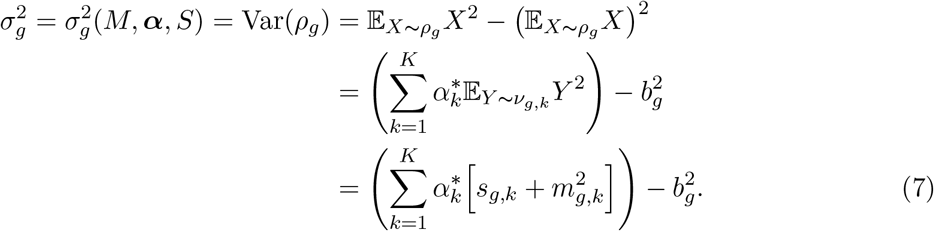

The likelihood of observing data 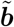 then follows straightforwardly from the central limit theorem:

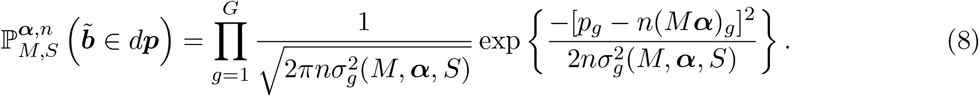

### Accounting for uncertainty in the design matrix

The above assumes exact knowledge of the individual distributions *ν_g,k_* (or rather their expectations *m_g,k_*), which is implausible in experimental settings. Instead, *M* needs to be estimated from data through some estimator 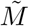, which we conveniently take to be the sample mean of expression across cells within each cell type, 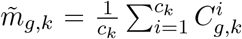, where 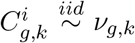, and *c_k_* denotes the number of single cells of cell type *k*. With this additional correction, (5) becomes

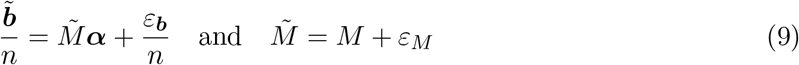

where *ε_M_* is a matrix of entries (*ε_M_*)*_g,k_* independently following 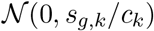 distributions. The second major difference between RNA-Sieve and existing tools (especially those based on least-squares methods) is the correction of the least-squares-type likelihood (8) by this stochasticity in the design matrix:

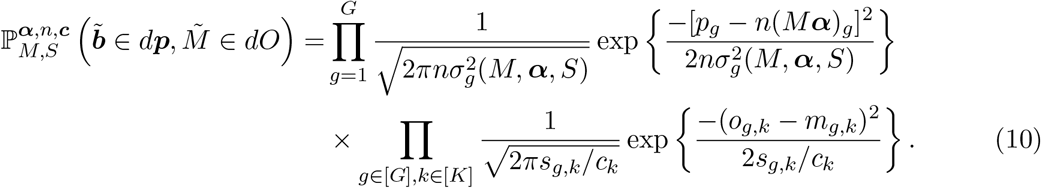

Our method utilizes the likelihood shown in (10), the suitability of which depends on a few implicit assumptions that are worth examining. The first is that the large number of cells assayed in an experiment permits us to use asymptotic theory and apply the classical CLT. As a result, we can write down a likelihood for our observations using normal distributions as long as Var(*ν_g,k_*) < ∞, which is true since gene expression profiles are necessarily bounded. Secondly, we suppose that the errors arising from estimating ***b*** and *M* are independent. This is appropriate as the bulk and single-cell experiments are performed separately. We additionally presume that expression levels in different genes are independent, as are those in different cells. It is unclear whether the latter is completely true in practice, though there is little evidence to the contrary. On the other hand, expression levels across genes within samples (either bulk or individual cells) are liable to be somewhat dependent due to expression co-regulation and the nature of the sampling process performed in RNA-seq. Given the large number of genes assayed, the latter co-dependence is apt to be fairly small. Meanwhile, co-expression estimation in single cells remains an open problem independent of deconvolution tasks, and so is not accounted for in RNA-Sieve. Once correlation structure is known however, it is straightforwardly incorporated into the likelihood we propose.

### Joint deconvolution of multiple bulk samples

If it is known that multiple bulk gene expression vectors share the same constituent cell type expression profiles, we can gain statistical strength and decrease the computational burden by inferring their mixture proportions jointly rather than individually. Assuming statistical independence of the bulk sample observations, we must simply augment the likelihood in (10) by including the *N* − 1 additional mixtures in *A* = (*α*_1_, …, *α_N_*) ∈ ℝ^*K × N*^, 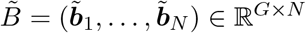 and ***n*** = (*n*_1_*, …, n_N_*):

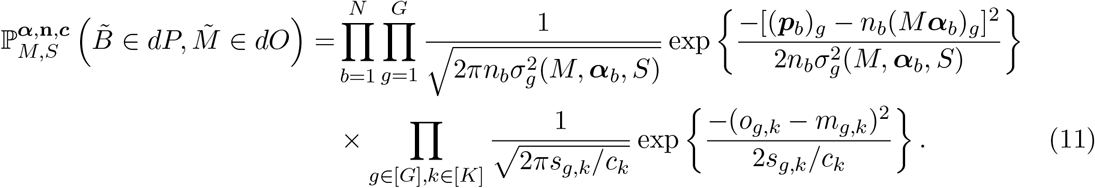

This increase in power depends solely on the statistical independence of distinct bulk samples rather than their respective cell type compositions. In fact, for the purposes of denoising the reference *M*, samples of dissimilar compositions are preferable because they provide non-redundant information. Conversely, bulk samples exhibiting heterogeneity in gene expression patterns (e.g., through differential expression) without corresponding reference matrices *M* amount to model misspecification, and thus may negatively impact inference. This impediment is information-theoretically unavoidable and therefore a challenge for all deconvolution methods. In our particular applications we did not find a strong effect of sample heterogeneity on our results; for instance, simultaneous deconvolution with mice of different ages yielded highly similar results to when we stratified by age. In the case of cell types with strong expression differences across different phenotypes, this may not hold, however.

### Data Pre-processing Procedure

Due to the well-known influence of technical variability in scRNA-seq data, we suggest that users of RNA-Sieve perform their own quality control filtering of cells and genes prior to running our software in addition to their preferred normalization. Given the potential complexity of these patterns in general, we feel that manual cleaning is more reliable than automated procedures. Nonetheless, we implement a simple, largely optional, cell filtering and normalization scheme to ensure the accuracy of results when the user has chosen not to perform their own quality control. Our procedure attempts to do the following:

1. Remove low-quality cells with anomalously low or high library sizes (≥ 3 median absolute deviations away from the median value of total number of reads per cell in each cell type)
2. Normalize read counts in cells (re-scale reads so that all cells have the median number of reads from across all cells);
3. Identify and remove cells which may be mislabeled or are simply extremely different from other cells with the same cell type label (≥ 3 median absolute deviations away from the median value of inter-cellular pairwise distances in each cell type)
4. Identify and retain genes which are expressed sufficiently often (≥ 20% non-zero measurements in at least one cell type).

We note that the first three steps are optional whereas step 4 is necessary to remove lowly expressed genes, whose presence may result in poor optimization outcomes due to creating biologically implausible expressions (a non-zero bulk expression can never be realized as a convex combination of zero or almost zero, low variance, single cell expressions).

We implemented two additional layers of gene filtering which we found improved robustness to cross-protocol differences in reference and bulk gene expression measurements. The motivation behind these steps is as follows:

1. By virtue of being a convex combination of expression levels from different cell types (under our generative model (10)), a gene’s *true* expression *b_g_* must necessarily lie between its smallest and largest corresponding expressions *m_g,k_* across cell types *k* ∈ [*K*]. That is,

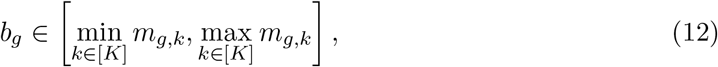

which naturally motivates a filtering scheme based on violations of these constraints. Of course, these inequalities do not necessarily hold in the presence of observational noise, which may push a gene’s bulk expression outside of its theoretical extremes. However, a stochastic version of (12) persists in that

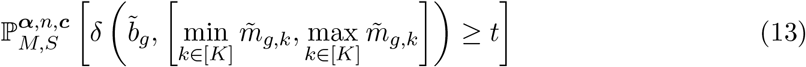

decays in *t* with sub-Gaussian tails (with constants depending on 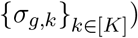, where *δ*(*p, A*) = inf*_a∈A_* |*p* − *a*| is the shortest distance of the point *p* to a set *A*. It is thus plausible to filter out all genes for which (13) is sufficiently small (in principle, computing the precise tail bounds (13) requires access to the true parameter ***α***, which prior to deconvolution is not available; however, reasonable upper bounds of (13) can be calculated independently of ***α***).
2. Gene expression profiles may experience (occasionally drastic) shifts when measured with distinct protocols. For example, mean and variance estimates of some gene expression levels may correlate little, or even not at all, across data generated using Smart-Seq2, UMI-based, or bulk RNA-seq technologies. To identify and remove these genes, we resort to a handful of empirically effective filtering steps. Specifically, we remove a gene if it presents as an outlier (as measured by median absolute deviations from the median) in any of the following summary statistics:

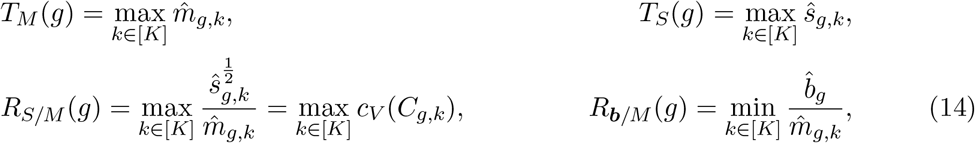

where *c_V_* (*C_g,k_*) denotes the coefficient of variation associated with expression profiles of gene *g* in cell type *k*. While the choice of these summary statistics was primarily guided by empirical considerations, they do reveal intuitively plausible and previously observed patterns: *T_M_, T_S_* and *RS/M* reflect the fact that severe over- or under-expression, or high degrees of variability in expression are not well-preserved across protocols, whereas 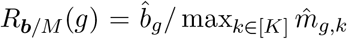 directly assesses any abnormal conversion factors between bulk and reference protocols.

In our experience, applying these filters based on (13) and (14) on top of the basic cell filter retains between 3, 000 − 12, 000 genes on which to perform the deconvolution task.

### Optimization and Estimation

We estimate ***α***, the cell type proportions for a given bulk sample, using the MLE which arises from maximizing (10). Given the number of free parameters (*GK + K* in total, corresponding to *M*, ***α***, and *n*) and structure of the likelihood, this is non-trivial, with standard optimization schemes commonly failing or returning sub-optimal solutions. On its face, the shape of (10) is reminiscent of loss functions appearing in so-called Total Least Squares formulations (see, e.g., (Golub and Van Loan, 1980)), whose minimizers can typically be found through SVD-decompositions. However, the presence of entry-wise uncertainties *ε_M_* and the dependence of *ε*_***b***_ on ***α*** render such spectral tools inapplicable to our setting; indeed, the corresponding linear algebraic problem consists of finding low-rank approximations to the concatenation of *M* and ***b*** in a Frobenius norm with ***α***-dependent weights, for which no satisfactory theory exists. We thus propose an alternating maximization scheme which iteratively estimates and updates ***α***, *M*, and *n* (and consequently 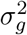) via a combination of quadratic programming and gradient descent. Despite the increased computational burden relative to W-NNLS or similar techniques, we find that convergence times remain reasonable, requiring between 15-40 minutes on typical data sets of 10, 000+ genes and 6 cell types using a modern laptop computer. We sketch an overview of our optimization procedure below in Algorithm 1 (where 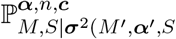 refers to (10) with ***σ***^2^(*M*, ***α****, S*) kept fixed at ***σ***^2^(*M* ^1^, ***α***^1^*, S*)).

Implementations of Algorithm 1 are currently available in Python and Mathematica. While these implementations differ slightly, they both agree on the following fundamental design choices:

1. Instead of maximizing 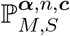, we minimize 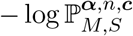, rendering lines 5 and 6 as quadratic programs which can be solved efficiently.
2. Lines 7 and 13 can be solved explicitly by differentiating (10) and finding the zeros of the resulting algebraic fractions in *n*. Thus, these steps do not require any explicit optimization scheme.
3. The optimizations in lines 11 through 13 proceed via gradient descent (or a variation thereof), and so could possibly require long runtimes. However, the coarser maximization (minimization, cf. item 1) in lines 5–7 typically improves the objective function to such an extent that only two or three more iterations are required. Moreover, both sets of optimizations are amenable to parallelization.
4. Algorithm 1 straightforwardly generalizes to the setting of jointly inferring mixture proportions in an arbitrary number *N* of bulk samples (cf. the remark in Mathematical Model). Both of our implementations support this generalized deconvolution.

Lastly, we note that although the alternating optimization in lines 5–7 is not guaranteed to converge, the second round of maximization in lines 11–13 is a proper coordinate descent and is therefore guaranteed to reach a local minimum.

**Algorithm 1:**
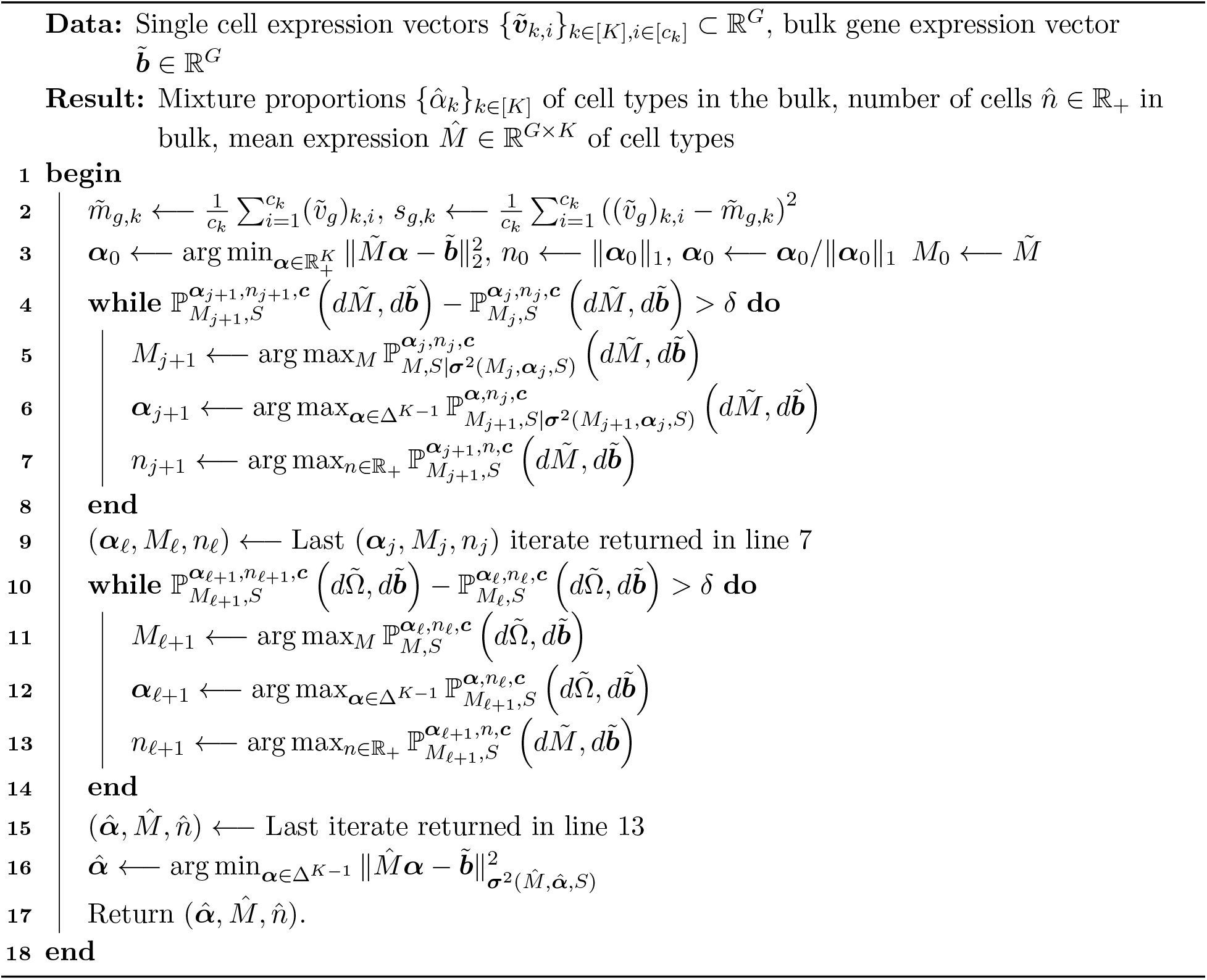
Find MLE of ***α***.

### Confidence Intervals

As indicated in Extension to Confidence Regions, the explicit generative modeling of (10) allows us to not only compute precise point estimators of ***α*** and *n*, but also to quantify this precision through confidence regions. More concretely, since our model is well-behaved in the sense of satisfying all assumptions in, say, Theorem 9.14 of Keener (2011), we expect our estimates 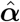 and 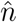 to be distributed normally around the true configuration ***α***\*, *n*\* with covariance matrix given by the inverse of the Fisher information 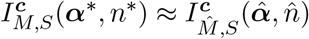. Given such asymptotic normality, it is straightforward to construct both marginal confidence intervals (from the diagonal entries of 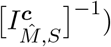 as well as *K*-dimensional confidence regions around 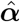. Generically, there are infinitely many possibilities for choosing such confidence regions from 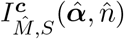, which we acknowledge by providing the entire (inverse) Fisher information to the user to allow computation of the confidence volume of their choosing. One option is tcoalculattehe canonical (that is, Lebesgue volume-minimizing) *q*-confidence region 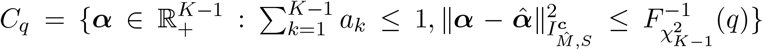, where ||***v***||_Σ_ = ⟨***v***, Σ^−1^***v***⟩ is the Mahalanobis norm of ***v*** associated with covariance matrix Σ, 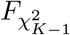, denotes the CDF of a *χ*^2^ variable with *K* − 1 degrees of freedom, and where we reparameterize ***α*** in order to account for the simplex constraint in our computation of the Fisher information matrix; this option is included as the default.

Confidence intervals derived in this manner are, as a consequence of the aforementioned Theorem 9.14 in Keener (2011), necessarily well-calibrated *if* data adhere to our generative model (10). As we have observed in Data Pre-processing Procedure, this may not hold when the use of different protocols in the reference and bulk experiments induce significant distributional shifts. Nonetheless, we can still provide conservative, yet well-calibrated, confidence regions by generalizing the Fisher information 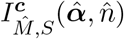 to the Godambe information matrix 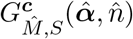 of the data (Godambe, 1960). If protocol mismatches result in the true generating distribution ℚ of the data not lying within our model family 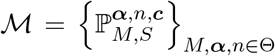, where Θ ⊂ ℝ^*G*·*K*+(*K*−1)+1^ is the space of all possible parameter configurations, then 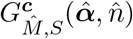 de-scribes the Gaussian fluctuations of 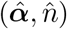 around the relative-entropy-projection of ℚ onto 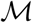; i.e., 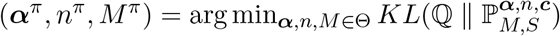, where *KL*(*ν || μ*) denotes the relative entropy (also known as Kullback-Leibler divergence) between two probability distributions *ν* and *μ*. Thus confidence regions for 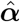 based on 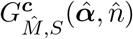 are still well-calibrated, assuming that ***α***\* ≈ ***α****^π^*, which is plausible given that distributional shifts induced by protocol differences appear to affect expression means (the entries of *M*) primarily through global scaling. In the absence of distributional mismatches, the Godambe information matrix 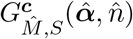 automatically collapses to the (empirical) Fisher information matrix 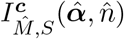, and so our confidence region estimation proceeds through 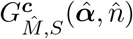 in both within- and cross-protocol settings by default, though can be adjusted manually if so desired. Occasionally, especially when constituent cell types are closely related to each other, the resulting covariance matrices may be nearly singular, in which case their inversion poses computational difficulties. To sidestep potential numerical instabilities, we subsample genes according to their incurred residual values. This reduces the probability of collinearities and produces more well-behaved confidence intervals. Simulations both across and within protocols confirm the utility of our confidence regions, and therefore the validity of the ***α****^π^* ≈ ***α***\* assumption, assessed in this manner (cf. Figure 7). Namely, our estimated 95%-confidence intervals contain the true cell type proportions 96.7% and 91.8% of the time in the *Tabula Muris Senis* pseudobulks, 95.8% of the time in the Monaco et al. bulk samples, and 90.3% of the time in the Newman et al bulk samples.

### A Note on *n*

One of the parameters inferred by our model is *n*, the number of cells in the bulk sample. This parameter is accurate and physically meaningful in within-protocol deconvolutions, or cross-protocol experiments where relative amplification factors are explicitly known. However, it loses interpretability when the relative scales across protocols are unclear, and so we sought to verify that both our inferred proportions and computed confidence intervals are robust even in such situations. Numerical experiments in which we re-scaled the bulk samples to artificially manipulate the inference of *n* showed no degradation in performance over a wide range of values, providing additional support beyond the observed high-quality results in both *in silico* and real bulk cross-protocol deconvolutions (Figure S10). Moreover, theoretical computations suggest a fairly weak dependence of confidence intervals on *n*, which are instead driven primarily by the total number of genes available for deconvolution.

### Comparison of runtimes

We found that all considered deconvolution algorithms could be successfully run in no more than a few hours on a laptop computer for the data sets we considered. RNA-Sieve runtimes ranged from 15-40 minutes, as did those of Scaden. Because of the straightforward manner in which we construct the signature and variance matrices for cell types, RNA-Sieve’s runtime is not sensitive to the size of the scRNA-seq reference. This is not the case for DWLS, whose runtime we found grew quickly with the size of the data set due to the model fitting involved in its signature gene inference procedure. For most cases, DWLS runtimes were also in the 15-40 minute range, but for some of the larger single-cell reference panels with many cell types, the runtime could extend to a few hours. CIBERSORTx typically ran in 5-15 minutes. The remaining methods (SCDC, MuSiC, Bisque, and NNLS) were quite fast, with runtimes of no more than a couple of minutes, though SCDC and MuSiC may take a few extra minutes if their tree-based deconvolution modes are used.

### Benchmarking procedures

We employed two distinct approaches to benchmark computational deconvolution methods. The first, the construction of so-called “pseudobulk” experiments, is a common strategy which aggregates scRNA-seq measurements across cells in order to construct gene expression mixtures which we treat as bulk samples with known ground-truth cell type proportions. For this task, we used data from the *Tabula Muris Senis* Consortium which covers many organs/tissues and cell types in two different single-cell experimental protocols–Smart-Seq2 and 10x Genomics Chromium. Specifically, we utilized bladder, kidney, large intestine, limb muscle, liver, lung, mammary gland, marrow, pancreas, skin, thymus, tongue, and trachea in these *in silico* experiments (see Table S2). For each tissue, four different deconvolutions were performed. For cross-protocol deconvolutions, one in which the reference came from Smart-Seq2 data with 10x Chromium pseudobulk and one in the reverse configuration. For within-protocol deconvolutions, the reference and pseudobulk were built using (non-overlapping) cells from the same protocol. For all pseudobulk deconvolution scenarios, a single reference set and pseudobulk was constructed. All eligible cells from each protocol were used. For scRNA-seq data from *Tabula Muris Senis*, the cell filtering procedure described in Data Pre-processing Procedure was applied.

Our second approach exploited the availability of bulk RNA-seq data sets with knowledge about true cell type proportions. For the PBMC and neutrophil scRNA-seq data sets, cells were filtered after manual inspection. Due to the large number of neutrophils in the available data set, 250 cells were randomly sampled from one individual for use with the Newman et al. reference and 1250 across three individuals were randomly sampled for use with the 10x Genomics reference. We considered four different scenarios:

1. Breast cancer and fibroblast cell lines and mixture from Dong et al. (2020);
2. Reference PBMCs and neutrophils from Newman et al. (2019) and Xie et al. (2020), respectively, with bulk whole blood from Newman et al. (2019);
3. Reference PBMCs and neutrophils from 10x Genomics and Xie et al. (2020), respectively, with bulk whole blood from Monaco et al. (2019);
4. Pancreatic islets from Xin et al. (2016) and Fadista et al. (2014).

The same data were used for all algorithms in each deconvolution, and all were run as described in their respective tutorials using default settings unless otherwise noted. When MuSiC was run, NNLS results were taken from MuSiC’s implementation; otherwise, the DWLS implementation was used. The corresponding scRNA-seq and bulk RNA-seq data files will be available at the Song Lab GitHub repository: https://github.com/songlab-cal/rna-sieve.

We chose to utilize the *L*_1_, *L*_2_, and *L*_∞_ distances, in addition to the Kullback-Leibler (KL) divergence, as our performance metrics for their ease of interpretation and ability to capture different aspects of algorithm performance. While the *L*_1_ and *L*_2_ distances, which we further average across cell types, are related to common notions of error such as the mean absolute deviation and root mean square error, the *L*_∞_ distance measures the largest difference between true and inferred values across all cell types and quantifies the worst-case performance in a deconvolution task. The KL divergence is a popular manner by which to compare probability distributions and so fits nicely with the deconvolution setting in which cell type proportions can be thought of as the sampling probability for an individual cell. It is also more sensitive to rarer cell types than the other considered metrics. We specifically chose to compute 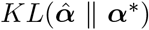 because it corresponds to the false positive rate when testing *H*_0_: ***α = α***\* against 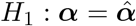 through a likelihood ratio test, and so is more relevant than 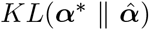.

## Data Availability

All data used in this paper are publicly available and are described in Table S4, with accession numbers included.

## Software Availability

RNA-Sieve is implemented in both Python and Mathematica. The source code is publicly available from the Song Lab GitHub repository: https://github.com/songlab-cal/rna-sieve.

## Acknowledgments

This research is supported in part by grant number R35-GM134922 from NIH and grant number CZF2019-002449 from the Chan Zuckerberg Initiative Foundation. Y.S.S. is a Chan Zuckerberg Biohub Investigator.

