## Supplemental Material for "A likelihood-based deconvolution of bulk gene expression data using single-cell references"

This file contains:

- Supplementary Tables S1–S4
- Supplementary Figures S1–S10

(a) Smart-Seq2 reference and 10x Chromium pseudobulk

| | RNA-Sieve | Bisque | CIBERSORTx | DWLS | MuSiC | NNLS | Scaden | SCDC |
|---|---|---|---|---|---|---|---|---|
| Bladder | 0.081 | **0.047** | 0.082 | 0.072 | 0.106 | 0.378 | 0.099 | 0.113 |
| Kidney | 0.095 | 0.109 | **0.028** | 0.055 | 0.110 | 0.249 | 0.062 | 0.083 |
| Large intestine | 0.076 | 0.082 | 0.300 | 0.123 | 0.108 | 0.226 | **0.042** | 0.136 |
| Limb muscle | 0.199 | 0.108 | 0.037 | 0.039 | 0.199 | 0.310 | **0.030** | 0.144 |
| Liver | 0.137 | 0.129 | 0.030 | 0.054 | 0.139 | 0.340 | 0.076 | **0.027** |
| Lung | 0.056 | 0.078 | 0.071 | 0.064 | 0.056 | 0.149 | 0.057 | **0.029** |
| Mammary gland | **0.020** | 0.258 | 0.072 | 0.029 | 0.047 | 0.371 | 0.083 | 0.058 |
| Marrow | 0.061 | 0.101 | 0.071 | 0.073 | 0.070 | 0.166 | 0.072 | **0.049** |
| Pancreas | **0.011** | 0.029 | 0.050 | 0.030 | 0.067 | 0.130 | 0.059 | 0.067 |
| Skin | **0.019** | 0.270 | 0.048 | 0.123 | 0.098 | 0.462 | 0.182 | 0.128 |
| Thymus | **0.017** | 0.050 | 0.098 | 0.331 | 0.127 | 0.482 | 0.030 | 0.120 |
| Tongue | **0.016** | 0.289 | 0.068 | 0.293 | 0.047 | 0.448 | 0.217 | 0.017 |
| Trachea | 0.108 | **0.097** | 0.166 | 0.165 | 0.142 | 0.252 | 0.110 | 0.154 |

(b) 10x Chromium reference and Smart-Seq2 pseudobulk

| | RNA-Sieve | Bisque | CIBERSORTx | DWLS | MuSiC | NNLS | Scaden | SCDC |
|---|---|---|---|---|---|---|---|---|
| Bladder | **0.002** | 0.066 | 0.059 | 0.085 | 0.156 | 0.044 | 0.167 | 0.261 |
| Kidney | 0.082 | 0.045 | 0.036 | **0.028** | 0.113 | 0.173 | 0.044 | 0.046 |
| Large intestine | 0.117 | 0.158 | 0.089 | 0.152 | 0.186 | 0.448 | 0.066 | **0.007** |
| Limb muscle | 0.137 | 0.132 | 0.037 | **0.013** | 0.122 | 0.142 | 0.102 | 0.177 |
| Liver | 0.107 | 0.056 | **0.032** | 0.050 | 0.126 | 0.164 | 0.052 | 0.070 |
| Lung | 0.092 | 0.069 | 0.029 | **0.021** | 0.130 | 0.153 | 0.074 | 0.045 |
| Mammary gland | **0.009** | 0.244 | 0.062 | 0.013 | 0.160 | 0.228 | 0.196 | 0.157 |
| Marrow | **0.070** | 0.110 | 0.124 | 0.097 | 0.113 | 0.147 | 0.111 | 0.121 |
| Pancreas | 0.121 | 0.085 | 0.137 | 0.054 | **0.023** | 0.117 | 0.111 | 0.173 |
| Skin | **0.037** | 0.191 | 0.162 | 0.050 | 0.168 | 0.676 | 0.109 | 0.192 |
| Thymus | **0.002** | 0.110 | 0.114 | 0.208 | 0.298 | 0.317 | 0.036 | 0.275 |
| Tongue | **0.006** | 0.254 | 0.022 | 0.143 | 0.672 | 0.672 | 0.191 | 0.672 |
| Trachea | 0.092 | 0.098 | **0.067** | 0.080 | 0.166 | 0.151 | 0.105 | 0.153 |

Table S1: **Deconvolution errors for different algorithms in pseudobulk experiments.** Deconvolutions were performed using the specified methods in thirteen organs using both Smart-Seq2 and 10x Chromium data from the *Tabula Muris Senis* experiment. Presented errors show the $L_1$ distance between the ground truth and inferred values divided by the number of present cell types. These values correspond to Table 1 and Figure 2 of the main text.

| **Organs** | **# cell types** | **Cell types** |
|---|---|---|
| Bladder | 2 | bladder cell, bladder urothelial cell |
| Kidney | 7 | B cell, epithelial cell of proximal tubule, fenestrated cell, kidney collecting duct principal cell, kidney loop of Henle ascending limb epithelial cell, macrophage, T cell |
| Large intestine | 3 | enterocyte of epithelium of large intestine, epithelial cell of large intestine, intestinal crypt stem cell |
| Limb muscle | 6 | B cell, endothelial cell, macrophage, mesenchymal stem cell, skeletal muscle satellite cell, T cell |
| Liver | 5 | B cell, endothelial cell of hepatic sinusoid, hepatocyte, Kupffer cell, myeloid leukocyte |
| Lung | 12 | adventitial cell, B cell, bronchial smooth muscle cell, CD4+ $\alpha\beta$ T cell, CD8+ $\alpha\beta$ T cell, classical monocyte, fibroblast of lung, myeloid dendritic cell, neutrophil, natural killer cell, non-classical monocyte, vein endothelial cell |
| Mammary gland | 3 | basal cell, luminal epithelial cell of mammary gland, stromal cell |
| Marrow | 9 | granulocyte, granulocytopoietic cell, immature B cell, late pro-B cell, macrophage, megakaryocyte-erythroid progenitor cell, naive B cell, precursor B cell, promonocyte |
| Pancreas | 3 | pancreatic A cell, pancreatic B cell, pancreatic D cell |
| Skin | 2 | basal cell of epidermis, epidermal cell |
| Thymus | 2 | DN4 thymocyte, thymocyte |
| Tongue | 2 | basal cell of epidermis, keratinocyte |
| Trachea | 5 | basal epithelial cell of tracheobronchial tree, chondrocyte, endothelial cell, fibroblast, macrophage |

Table S2: **Cell types for each organ in pseudobulk experiments.** These were the cell types used in pseudobulk experiments with the *Tabula Muris Senis* data. The order in which they are listed here matches their order in any figures based off of these experiments.

| Data attribute | RNA-Sieve requirements |
|---|---|
| Cell counts | The asymptotic analysis of RNA-Sieve relies primarily on the Central Limit Theorem, and so any cell counts that allow its application are sufficient. Because most gene expression counts reasonably follow Poisson or negative binomial distributions, having at least 30 cells is typically sufficient for accurate approximations. Unusually skewed distributions may necessitate ∼100-400 cells. |
| Number of reference individuals | RNA-Sieve does not rely on the presence of multiple individuals in the reference and performs inference reliably with any number of individuals. If multiple reference individuals are available, RNA-Sieve simply operates on the pooled mean and variance matrices. We currently do not recommend mixing data from different experimental protocols in the reference. |
| Reference and bulk protocols | In the case of differences in the data due to protocol mismatch in the scRNA-seq reference and bulk samples, potential nonlinear distributional shifts may need to be accounted for (linear differences are absorbed into the inference of $n$, see the A NOTE ON $n$ section in the main manuscript). Empirically, we found such the largest driver of such nonlinear shifts to be differences in the rates of null inflation. In some cases, this is compensated for by increased sequencing depth. Thus, deeply sequenced libraries can be analyzed without further correction, while sparser data sets may benefit substantially from the filtering steps detailed in the DATA PREPROCESSING PROCEDURE section of the main manuscript. |
| Jointly deconvolving multiple bulks | If each cell type is expressed similarly across bulk samples (i.e., cells are not differentially expressed in different bulk samples), joint deconvolution is recommended as it increases statistical power regardless of any heterogeneity in mixture proportions. If cell types display differential expression (due to biological or technical reasons), model misspecification becomes a concern and inference results may depend on the nature of the misspecification. In such cases, it is advisable to deconvolve different bulk samples separately. |

Table S3: **Guidance on RNA-Sieve usage across diverse data sets.** RNA-Sieve's accuracy is based on a generative model operating in an asymptotic regime. The mild criteria outlined above guarantee that the data to be deconvolved behaves in accordance with this asymptotic generative model.

| **Source** | **Description** | **sc/bulk RNA-seq** | **Multi-subject** | **# bulks** | **Known truth** | **Location** |
|---|---|---|---|---|---|---|
| Tabula Muris Senis | Many mouse organs | Both | Yes | ∼40/organ | No | GSE13204 |
| Dong et al. (2020) | Human fibroblasts, cell lines | Both | No | 1 | Yes | GSE136148 |
| Newman et al. (2019) | Human PBMCs | scRNA-seq | No | – | – | GSE127471 |
| Newman et al. (2019) | Human PBMCs | bulk RNA-seq | Yes | 12 | Yes | GSE127813 |
| 10x Genomics data sets | Human PBMCs | scRNA-seq | Yes | – | – | See link below |
| Monaco et al. (2019) | Human PBMCs | bulk RNA-seq | Yes | 12 | Yes | GSE107011 |
| Xie et al. (2020) | Human neutrophils | scRNA-seq | Yes | – | – | GSE137540 |
| Xin et al. (2016) | Human pancreatic islets | scRNA-seq | Yes | – | – | GSE81608 |
| Fadista et al. (2014) | Human pancreatic islets | bulk RNA-seq | Yes | 77 | No | GSE50244 |

Table S4: **Descriptions of data sets used.** Source–original publisher of data; description–species and organs/tissues assayed; sc/bulk RNA-seq–which protocol(s) were used to assay expression; Muti-subject–whether more than one individual was sampled in the data set; #*bulks*–the number of bulk samples, if applicable; Known truth–whether the true cell type proportions were known or experimentally estimated for bulk samples; Location–accession number where data sets can be found. For the PBMC data from 10x Genomics, we used "3k PBMCs from a healthy donor" and "4k PBMCs from a healthy donor" accessed at `https://support.10xgenomics.com/single-cell-gene-expression/datasets`.

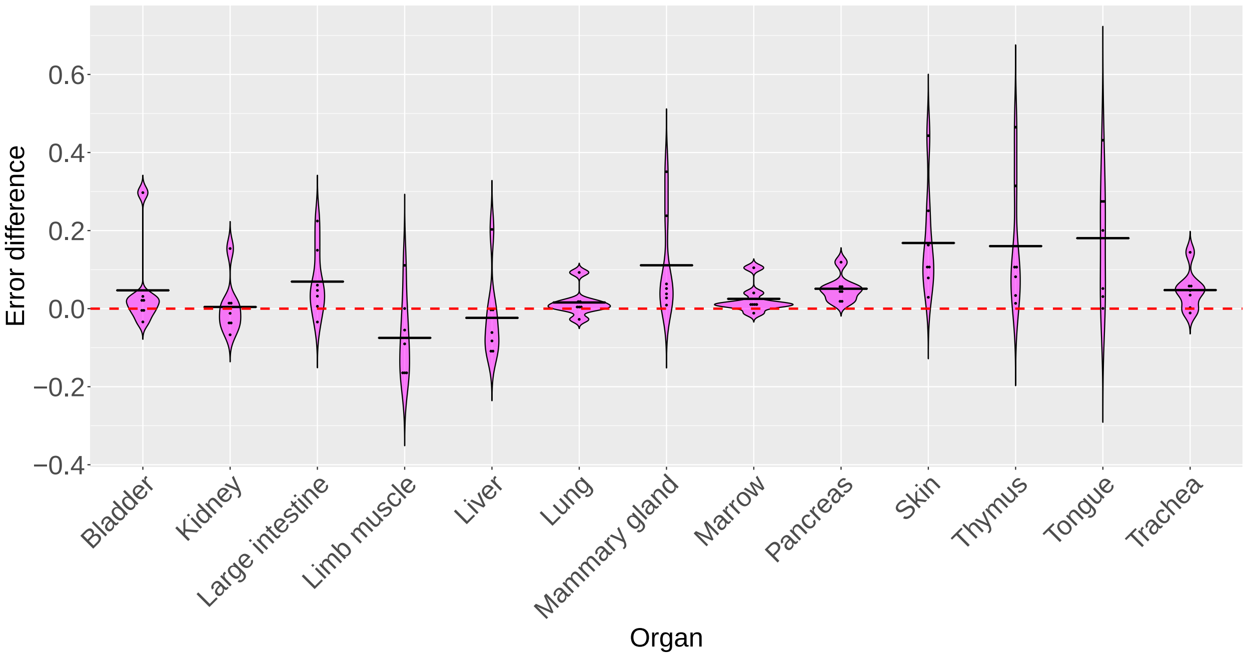


(a) **Smart-Seq2 reference and 10x Chromium pseudobulk**

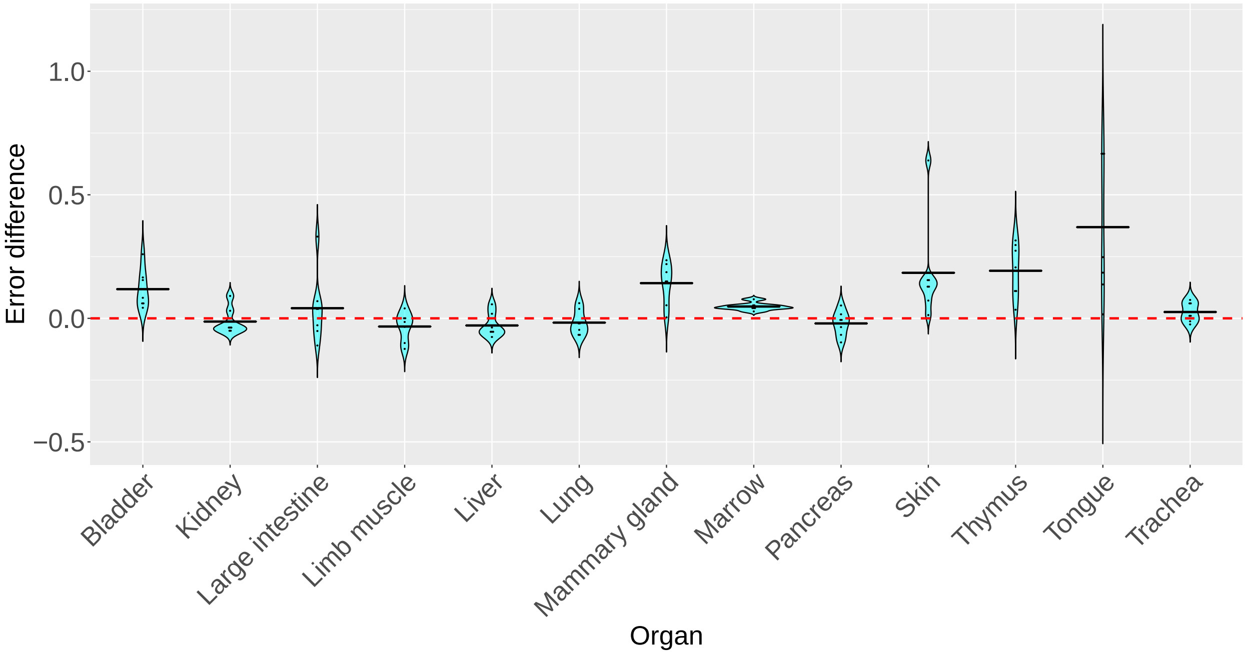


(b) **10x Chromium reference and Smart-Seq2 pseudobulk**

Figure S1: **Comparison of other methods to RNA-Sieve across 13 murine organs.** Pseudobulk experiments were performed in 13 different organs using data from the *Tabula Muris Senis* experiment. Errors were computed as the average $L_1$ error across cell types in each organ. For each organ, the difference in errors was computed between other methods and RNA-Sieve. Horizontal black bars correspond to the mean difference in error, and positive values indicate better comparative performance for RNA-Sieve.

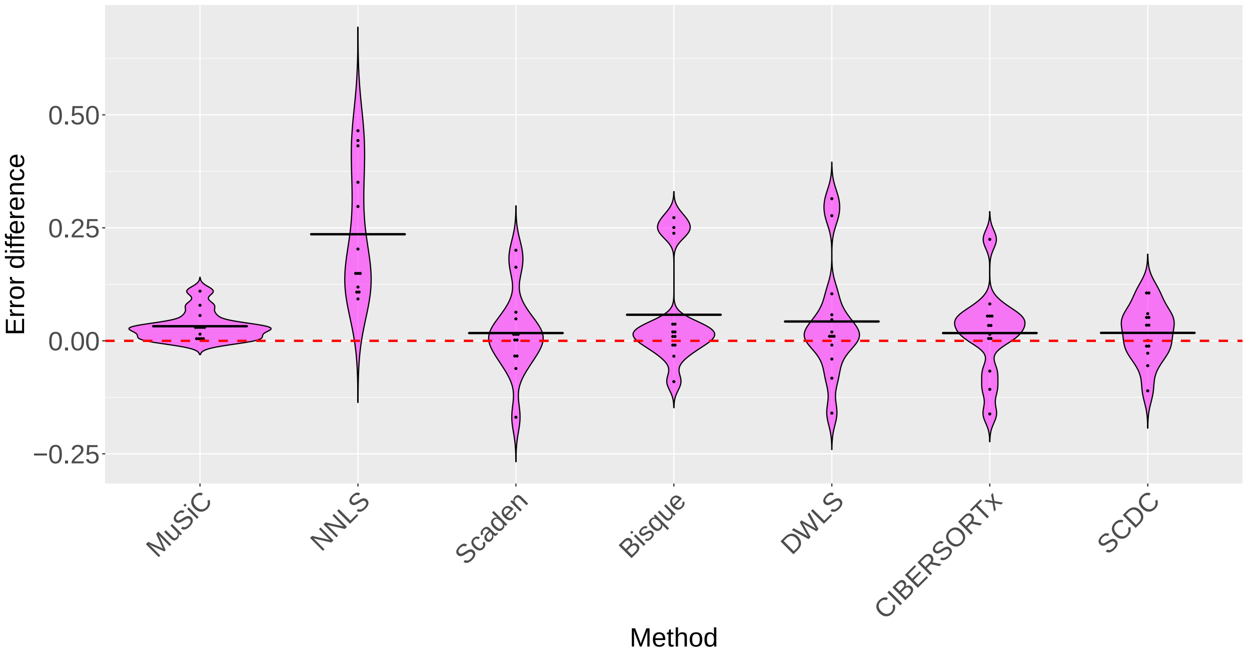


(a) **Smart-Seq2 reference and 10x Chromium pseudobulk**

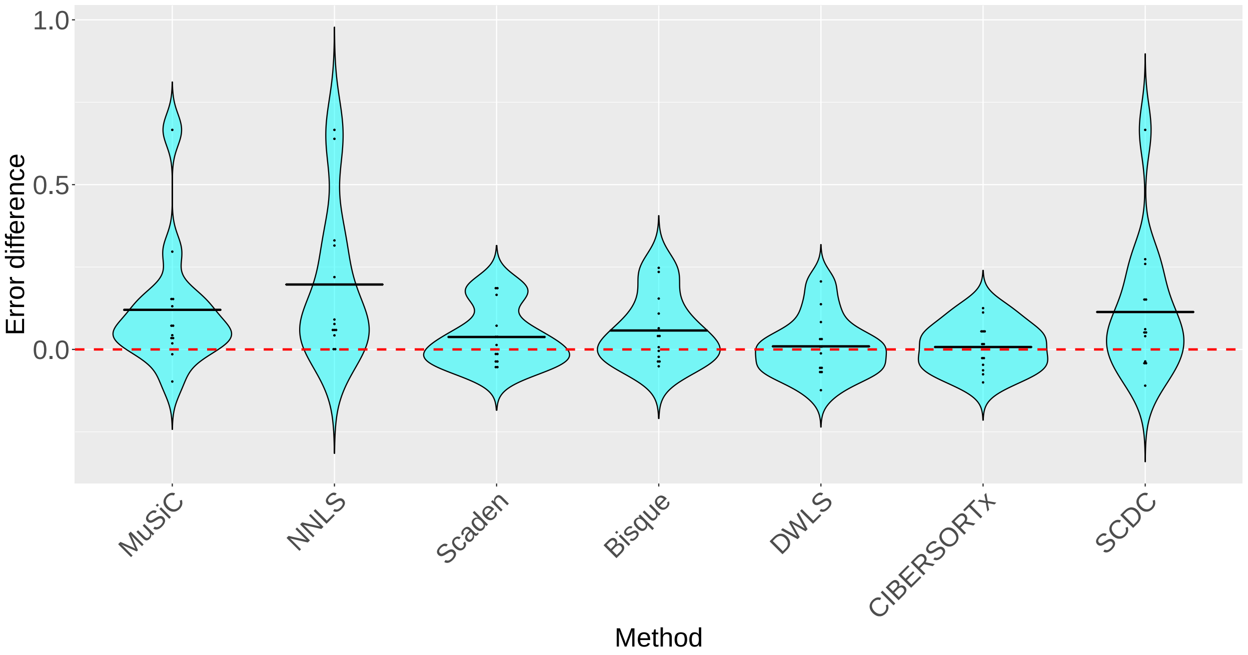


(b) **10x Chromium reference and Smart-Seq2 pseudobulk**

Figure S2: **Direct comparison of other methods to RNA-Sieve** Pseudobulk experiments were performed in 13 different organs using data from the *Tabula Muris Senis* experiment. Errors were computed as the average $L_1$ error across cell types in each organ. For each method, the difference in errors was computed between it and RNA-Sieve across each of the 13 organs. Horizontal black bars correspond to the mean difference in error, and positive values indicate better comparative performance for RNA-Sieve.

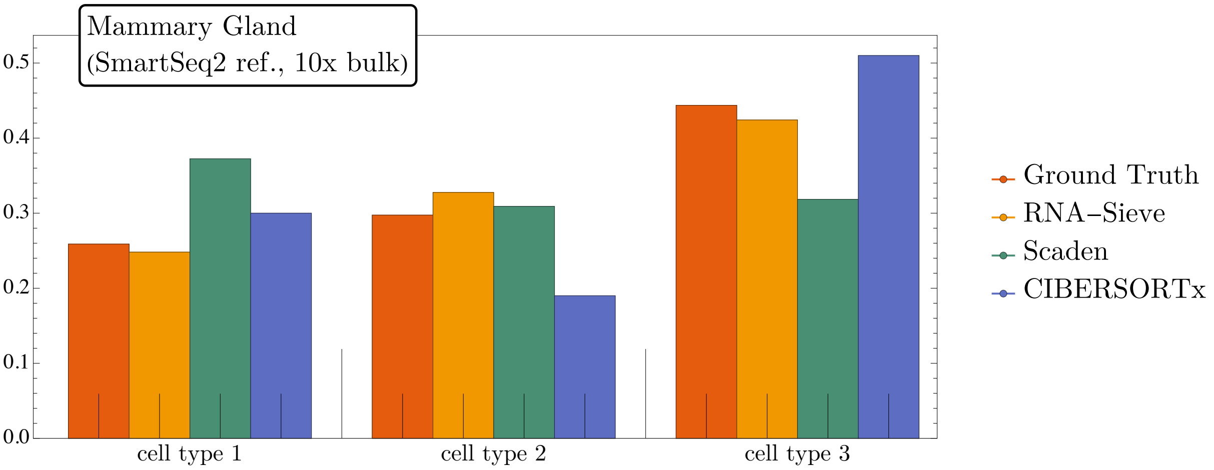


Figure S3: **Minor per-cell-type differences may result in major individual-cell-type deviations.** The average improvement of RNA-Sieve artificially appears minor because of our chosen error metric (average deviation from the true values) and averaging across cell types. This can be seen in the (real) example of deconvolving a 10x mammmary gland bulk from a SmartSeq2 reference in which RNA-Sieve (0.02), Scaden (0.08), and CIBERSORTx (0.07) may appear to perform similarly when only the raw error values are compared. However, closer inspection reveals that Scaden and CIBERSORTx exhibit large errors for some cell types whereas RNA-Sieve does not.

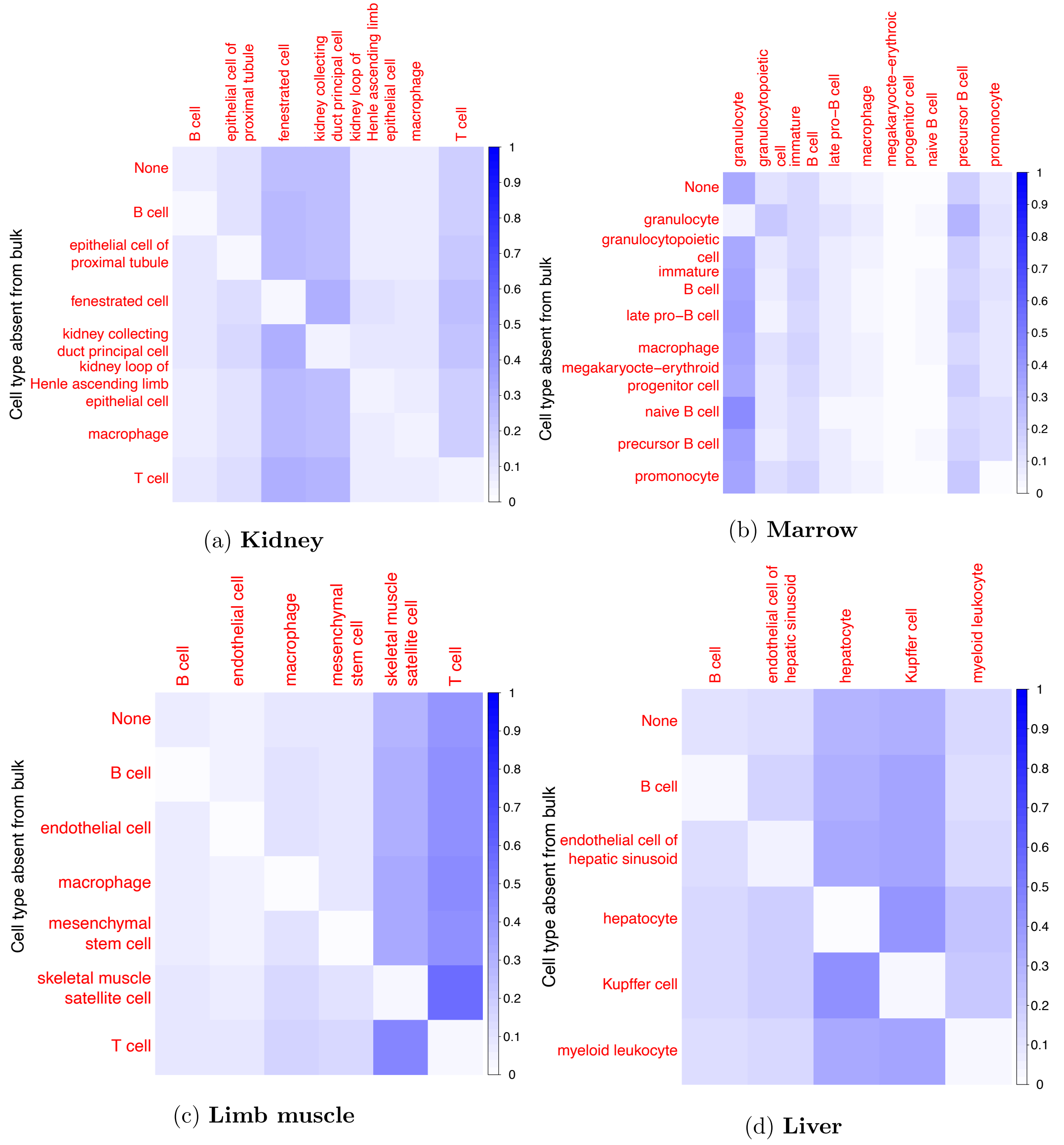


(a) **Kidney**

(b) **Marrow**

(c) **Limb muscle**

(d) **Liver**

Figure S4: **Deconvolution with extra cell types in the reference matrix.** Deconvolution was performed in pseudobulk experiments in four different organs. For each organ, we followed a leave-one-out procedure in which one cell type is removed from the pseudobulk at a time. Deconvolution was then performed with this extra cell type in the reference in order to examine RNA-Sieve's specificity. The top row shows the inferred proportions with no extra reference cell types. Darker colors indicate a higher estimated proportion value. Here we used 10x Chromium data for the reference and Smart-Seq2 for the pseudobulk.

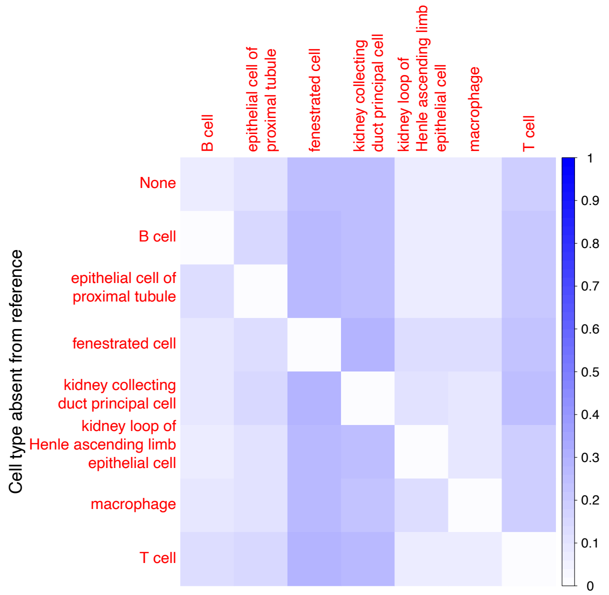


(a) **Kidney**

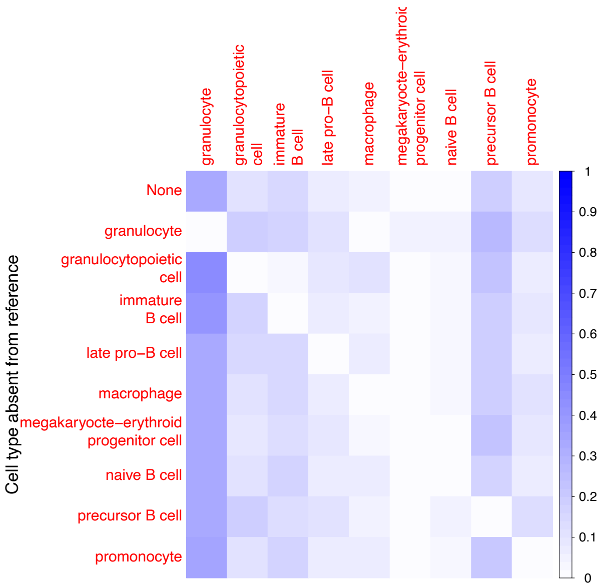


(b) **Marrow**

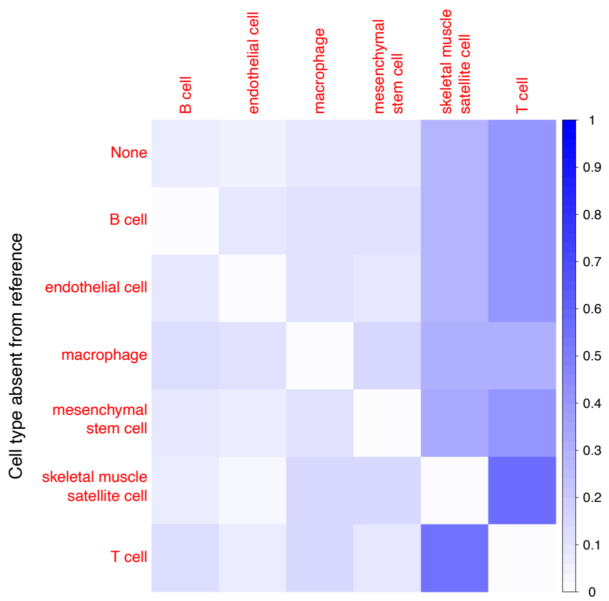


(c) **Limb muscle**

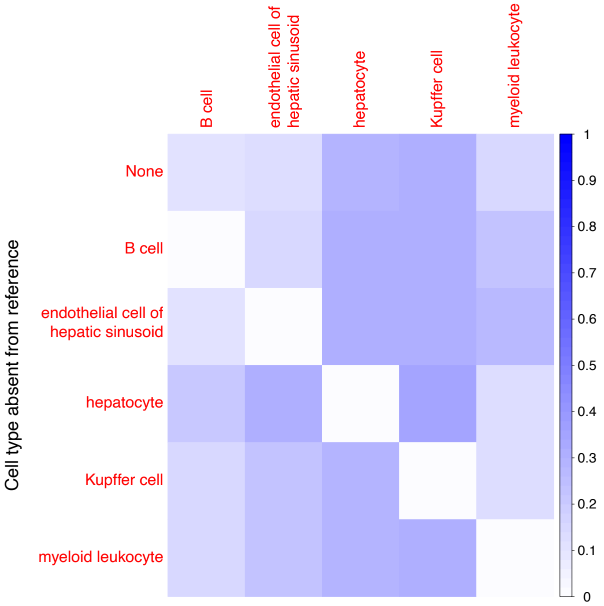


(d) **Liver**

Figure S5: **Deconvolution with missing cell types in the reference matrix.** Deconvolution was performed in pseudobulk experiments in four different organs. For each organ, we followed a leave-one-out procedure in which one cell type is removed from the reference at a time. Deconvolution was then performed with an extra cell type in the pseudobulk in order to examine RNA-Sieve's ability to handle such a misspecification. The top row shows the inferred proportions with no missing reference cell types. Darker colors indicate a higher estimated proportion value. Here we used 10x Chromium data for the reference and Smart-Seq2 for the pseudobulk.

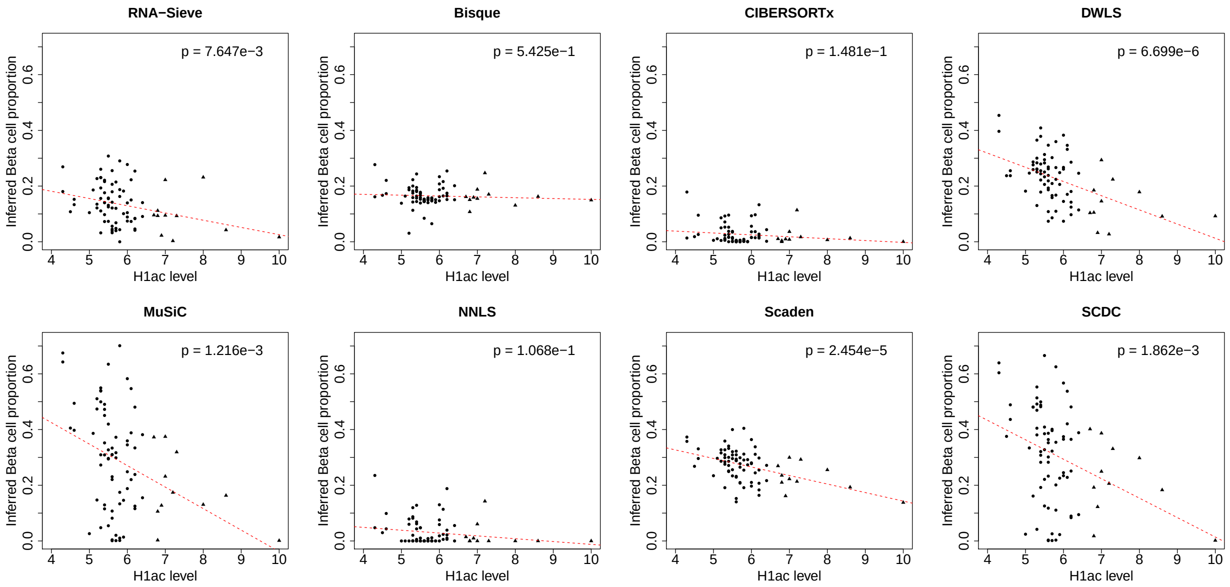


Figure S6: **Deconvolution results on validation data.** Single-cell expression data in pancreatic islets from Xin et al. (2016) was used as reference to deconvolve bulk RNA-seq data from Fadista et al. (2014). Each point represents the estimated beta pancreatic islet cell proportion one of 77 bulks with recorded HbA1c levels. The $p$-value is for a univariate regression on the estimated proportions. Circles correspond to healthy samples while triangles represent samples from diabetic patients.

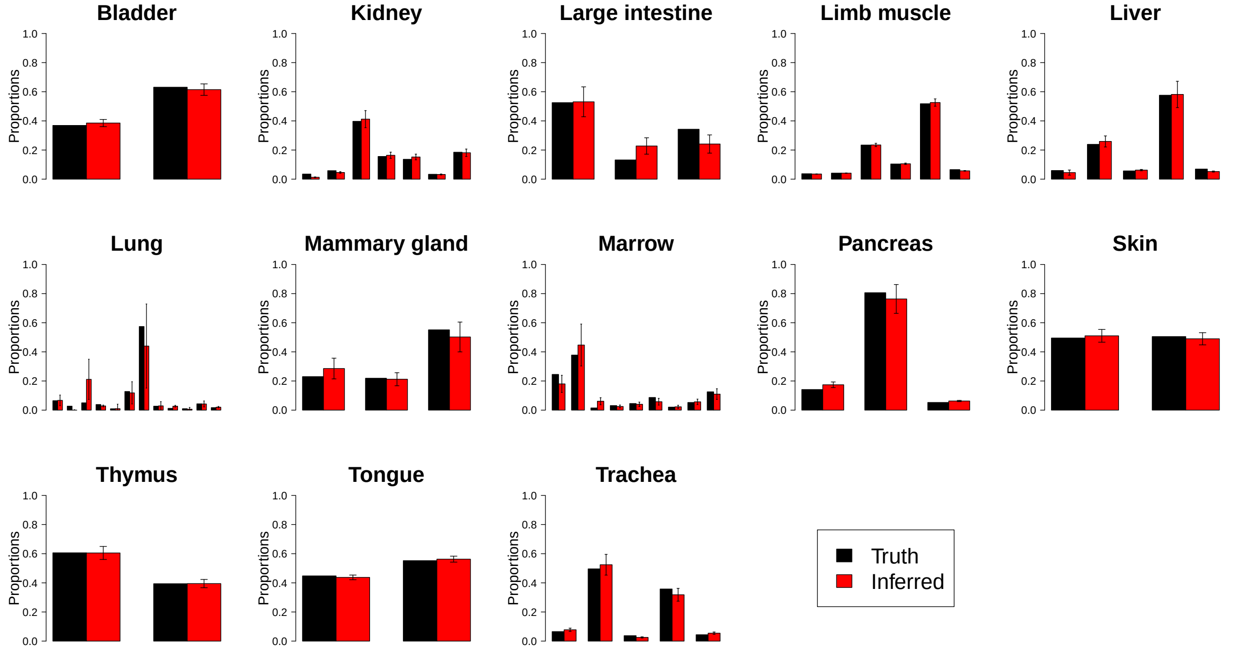


(a) **Within-protocol – 10x Chromium pseudobulk and reference**

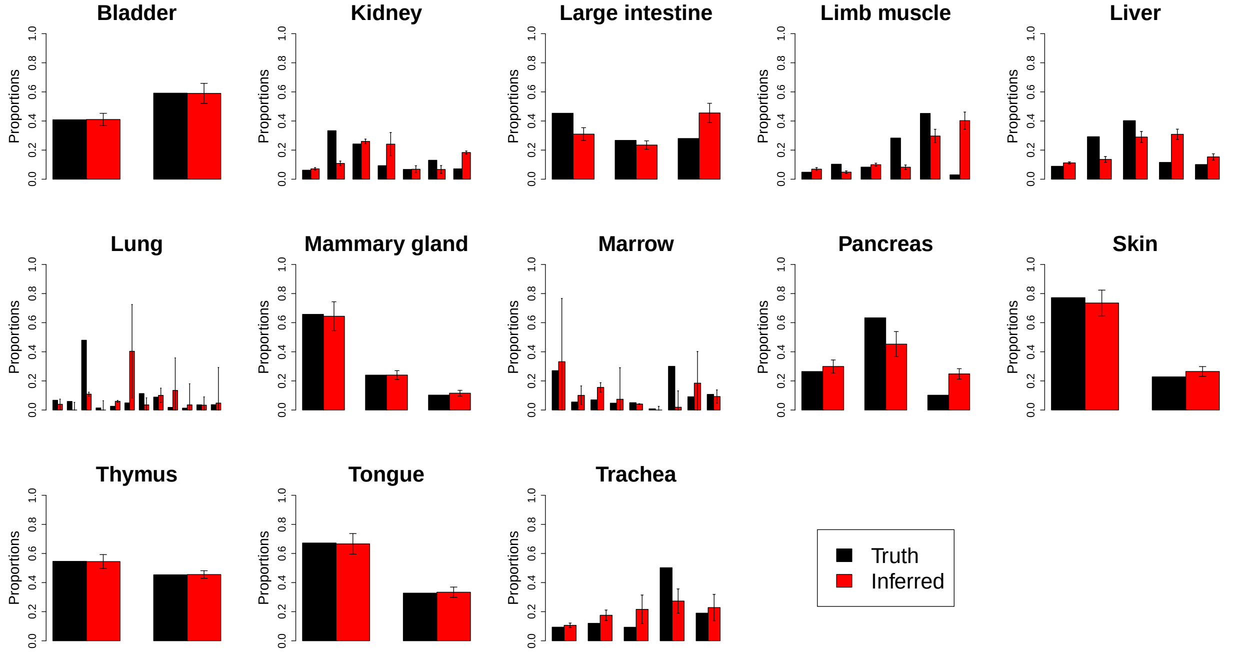


(b) **Cross-protocol – 10x Chromium reference and Smart-Seq2 pseudobulk**

Figure S7: **RNA-Sieve results with confidence intervals in pseudobulk experiments.** Inferred cell type proportions in pseudobulk experiments using data from the *Tabula Muris Senis* experiment. The black error bars on inferred proportions show the marginal 95% confidence intervals as computed from the estimated Fisher information produced by RNA-Sieve. Table S2 contains the cell types in each organ, which could not be displayed because of space constraints.

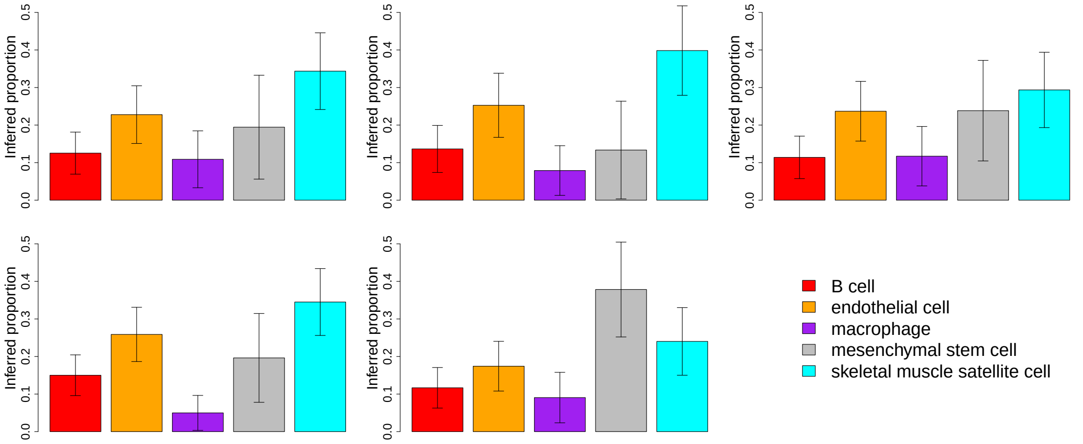


(a) **Limb muscle**

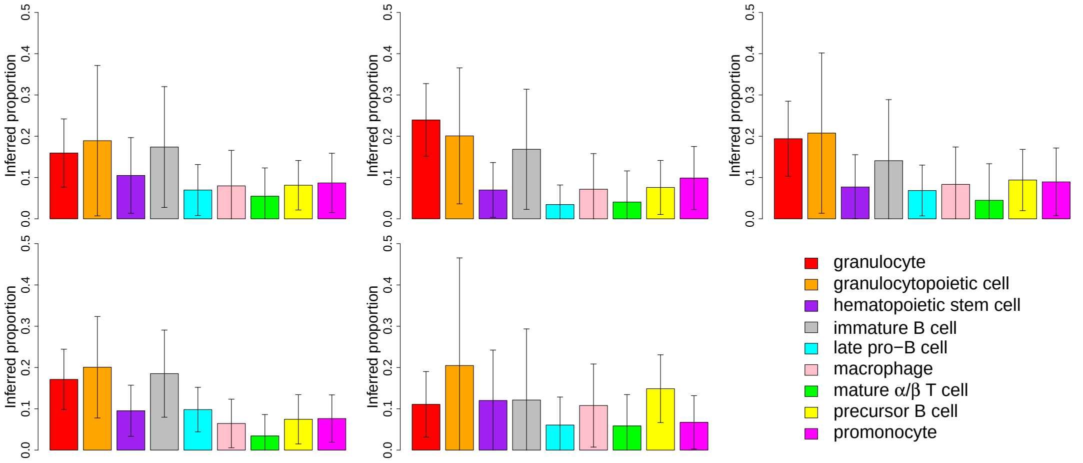


(b) **Marrow**

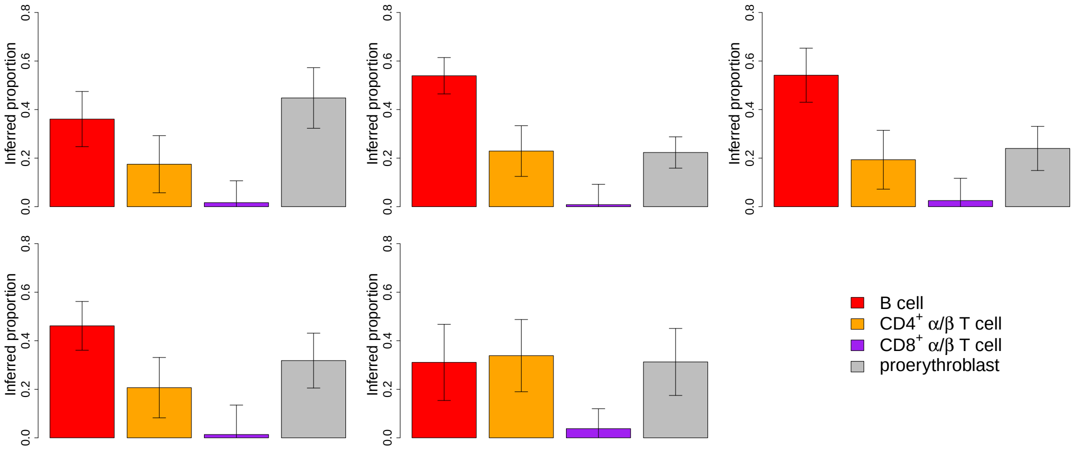


(c) **Spleen**

Figure S8: **Confidence intervals with real bulk samples** For all of each organ's samples, we produced estimated cell type proportions with 95% confidence intervals using RNA-Sieve. Smart-Seq2 data were used as the reference. Here we present five randomly chosen samples for each organ (out of ∼40); histograms showing the typical radii are displayed in Figure S9.

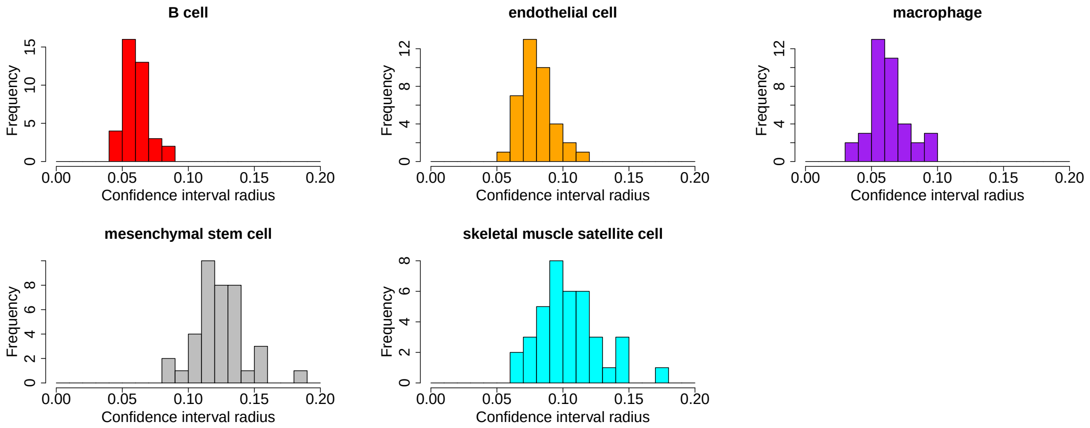


(a) **Limb muscle**

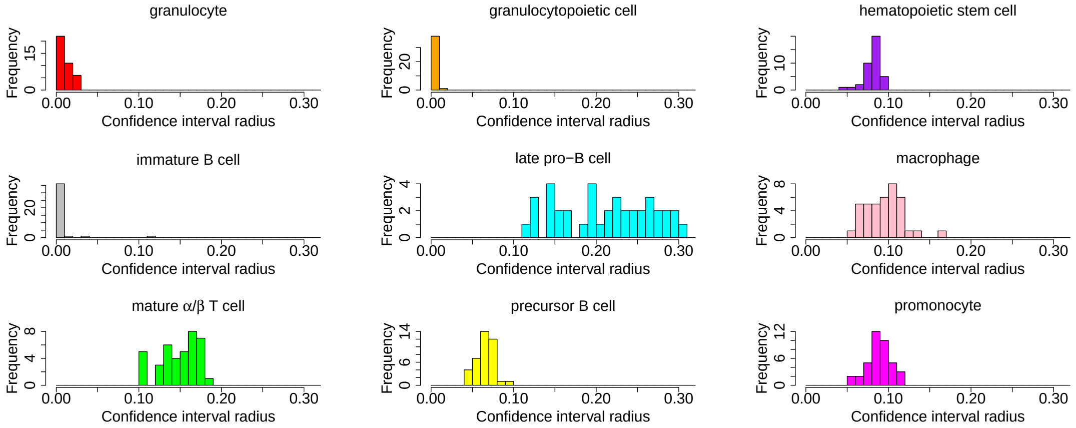


(b) **Marrow**

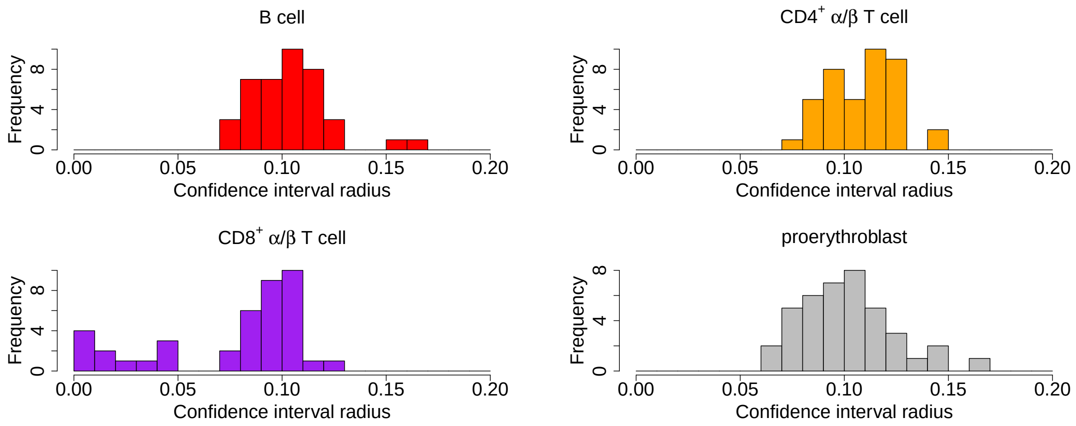


(c) **Spleen**

Figure S9: **Histograms of CI radii with real bulk samples.** The radius of the 95% confidence interval for inferred cell type proportions was computed using RNA-Sieve for each real bulk sample in the listed organs (∼40 per organ).

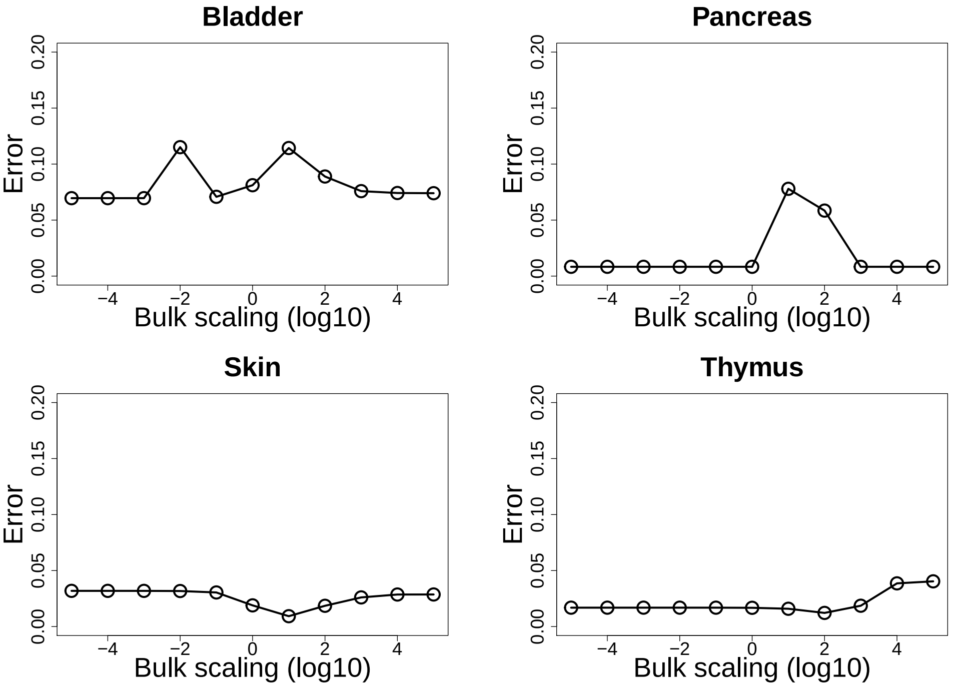


(a) **Smart-Seq 2 reference and 10x Chromium pseudobulk**

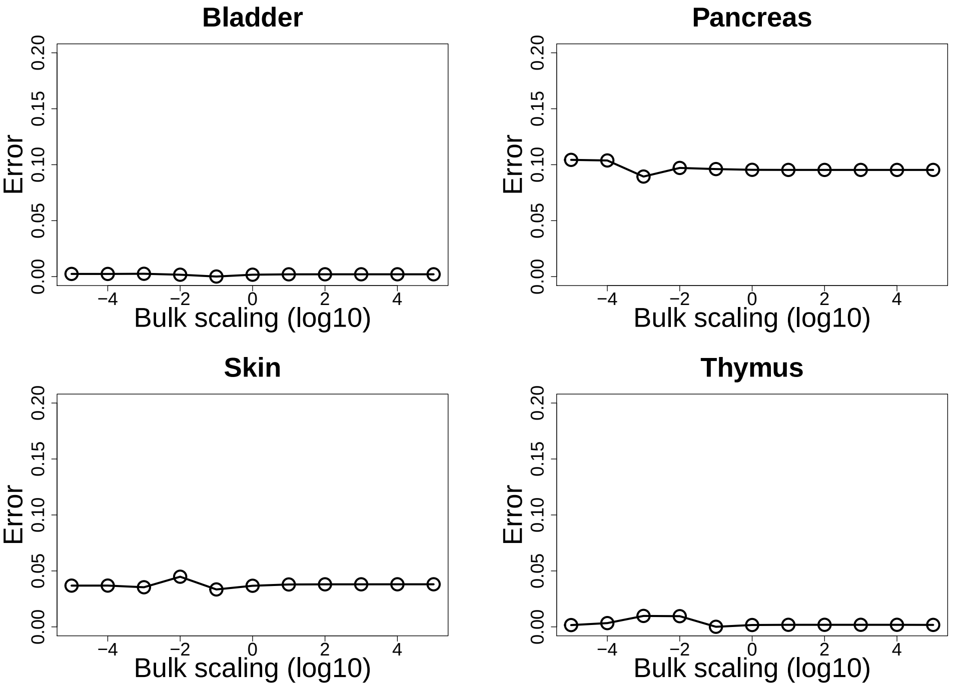


(b) **10x Chromium reference and Smart-Seq2 pseudobulk**

Figure S10: **Effects of different bulk scalings.** To determine RNA-Sieve's sensitivity to the the application of different scalings to bulk samples, we multiplied pseudobulk counts by different values across a wide range and performed deconvolution. We then computed RNA-Sieve's error for each value. Here we show representative results from four organs.
